# CREATE: cell-type-specific cis-regulatory elements identification via discrete embedding

**DOI:** 10.1101/2024.10.02.616391

**Authors:** Xuejian Cui, Qijin Yin, Zijing Gao, Zhen Li, Xiaoyang Chen, Shengquan Chen, Qiao Liu, Wanwen Zeng, Rui Jiang

## Abstract

Identifying cis-regulatory elements (CREs) within non-coding genomic regions—such as enhancers, silencers, promoters, and insulators—is pivotal for elucidating the intricate gene regulatory mechanisms underlying complex biological traits. The current prevalent sequence-based methods often focus on singular CRE types, limiting insights into cell-type-specific biological implications. Here, we introduce CREATE, a multimodal deep learning model based on the Vector Quantized Variational AutoEncoder framework, designed to extract discrete CRE embeddings and classify multiple CRE classes using genomic sequences, chromatin accessibility, and chromatin interaction data. CREATE excels in accurate CRE identification and exhibits strong effectiveness and robustness. We showcase CREATE’s capability in generating comprehensive CRE-specific feature spectrum, offering quantitative and interpretable insights into CRE specificity. By enabling large-scale prediction of CREs in specific cell types, CREATE facilitates the recognition of disease- or phenotype-related biological variabilities of CREs, thereby expanding our understanding of gene regulation landscapes.

## Introduction

Gene regulation is a fundamental biological process that orchestrates gene expression through a sophisticated network of interactions among biomolecules, including regulatory factors and cis-regulatory elements (CREs) located in non-coding genomic regions^1, 2^. CREs, such as silencers, enhancers, promoters, and insulators^3, 4^, typically locate in the chromatin accessible areas^5, 6^ and play crucial roles in modulating gene expression by interacting with target genes through chromatin loops^7, 8^ and other regulatory mechanisms. These characteristics are essential for controlling cell-type-specific gene expression patterns, which contribute to cellular diversity, tissue homeostasis, and the development of complex biological traits^9–11^. Consequently, identifying and characterizing cell-type-specific CREs is vital for advancing our understanding of gene regulation in normal physiology and disease states.

Silencers, enhancers, promoters, and insulators each have distinct roles in gene regulation. Silencers suppress gene transcription, enhancers boost transcriptional activity, promoters initiate transcription, and insulators act as boundary elements to regulate gene expression^3, 4^. Due to the restricted understanding of CRE-specific genetic signatures, identifying and validating CREs through biological experiments is cumbersome, time- and resource-consuming^9, 12^. Massive genomic and epigenomic data benefited from the rapid advancement of high-throughput sequencing technologies^13–16^, has provided valuable opportunities for identifying cell-type-specific CREs using computational methods. For example, DeepSEA is a convolutional neural network (CNN) model based on genomic sequences, which can simultaneously predict chromatin-profiling data such as transcriptional factors (TFs) binding sites, histone modification sites, and chromatin accessible regions^17^. DanQ is a hybrid deep neural network that merges convolutional and recurrent architectures, aimed at quantifying the non-coding function of DNA sequences^18^. Enhancer-Silencer transition (ES-transition) is a deep learning model based on CNN for identifying enhancers and silencers specific to cell types in the human genome, and has been utilized to uncover the unique phenomenon of enhancer-silencer transitions^19^. DeepICSH integrates DNA sequences with various epigenetic features including histone modifications, chromatin accessibility and TF binding to predict cell-type-specific silencers^20^.

However, recently innovated computational methods encounter many limitations and challenges. First, most current efforts are dedicated to the identification of a single type of CRE^20–23^, particularly enhancers. In the past few decades, there has been extensive research on enhancers^24–30^, while silencers, which generally share many properties with enhancers^31^, have received little attention. Numerous undiscovered CREs and uncharacterized chromatin regions suggest an urgent need for a comprehensive and scalable method of multi-class CRE identification. Second, mainstream methods prevalently extract information from DNA sequences to distinguish CREs^17–19^, overlooking the cell type specificity of CREs. Incorporating multi-omics data, including chromatin accessibility and chromatin interaction, for characterizing cell-type-specific CREs can provide valuable insights into gene regulatory mechanisms and cell heterogeneity. Third, deriving interpretable biological implications from conventional deep learning models remains challenging^32, 33^, hindering the meaningful large-scale identification of CREs and the understanding of model-related biological variabilities of CREs.

To bridge these gaps, we propose CREATE (Cis-Regulatory Elements identificAtion via discreTe Embedding), a novel CNN-based supervised learning model that leverages the Vector Quantized Variational AutoEncoder (VQ-VAE)^34–36^ framework. CREATE integrates genomic sequences with epigenetic features to offer a comprehensive approach for the identification and classification of multi-class CREs. The VQ-VAE framework is particularly suited for this task because it can distill genomic and epigenomic data into discrete CRE embeddings, capturing the nuanced differences between various CRE types. This discrete representation facilitates the generation of a CRE-specific feature spectrum, providing both quantitative and intuitive insights into CRE specificity. CREATE’s ability to integrate multi-omics data enables it to overcome the limitations of previous methods by offering a more holistic view of CRE functionality and their cell-type-specific roles. Furthermore, CREATE demonstrates superior performance in accurately identifying CREs and exhibits robustness across diverse input types and hyperparameters. Its capability to perform large-scale predictions of CREs in specific cell types and to uncover disease- or phenotype-related biological variabilities in CREs underscores its potential as a powerful tool for constructing a comprehensive CRE atlas. In summary, CREATE represents a significant advancement in the computational identification of CREs, and provides a robust foundation for future research in gene regulation and its implications for human health and disease.

## Results

### Overview of CREATE

CREATE is an advanced CNN model based on the VQ-VAE framework^34, 35^ to predict and classify multi-class CREs from multi-omics data. Taking as input the one-hot encoded genomic sequence, the vector representing the chromatin accessibility scores for that sequence, and the vector representing the chromatin interaction scores for that sequence, CREATE is specifically crafted to capture discrete CRE embeddings, providing a comprehensive and interpretable characterization of CREs (Fig. 1a and Methods).

**Fig. 1.**
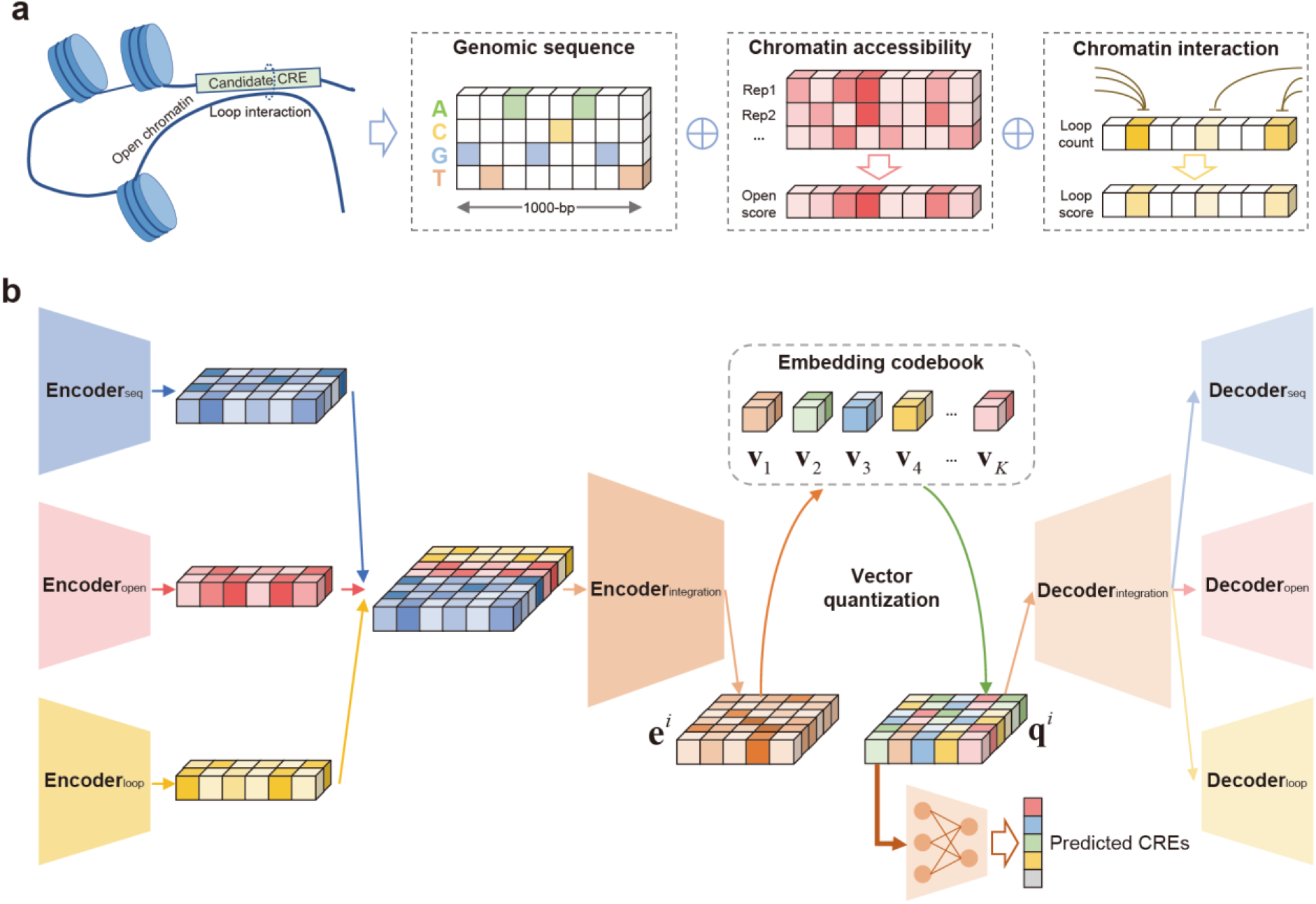
Overview of CREATE. **a**, The input of CREATE model. CREATE takes as input the genomic sequence, chromatin accessibility score and chromatin interaction score. **b**, The architecture of CREATE model. CREATE consists of encoders, a vector quantization module and decoders. The encoder module of CREATE combines encoders for multiple input-specific learning and an encoder for multiple input integration. For the *i*-th CRE, the encoder outputs the latent embedding 𝐞^𝑖^ of dimension *L’*×*D’*. By adopting split quantization, the latent embedding will be split into *L’*×*M* vectors 𝐞*_i,j_*^𝑖^ of dimension *D* and then quantized to 𝐪*_i,j_*^𝑖^ for the *i*-th CRE using embedding codebook with the size of *K*.

The architecture of CREATE includes (Fig. 1b and Methods): 1) Encoder Module: Each type of input data is initially processed by dedicated omics-specific encoders that transform the raw data into feature representations suitable for integration. Following this, the processed features are concatenated and passed through the integration encoder module, which synthesizes information from all input modalities to create a unified representation of the genomic context. 2) Vector Quantization Module: In this module, the output embeddings of encoder module are substituted with the closest counterpart in the discrete embedding space called “codebook”. In brief, the features in the codebook are concatenated to form the final CRE embeddings. Unlike traditional VAE-based models^32^ with fixed prior distributions, CREATE’s codebook is dynamic and updated during training. This flexibility allows the model to refine the discrete embeddings to better represent the underlying biological data. 3) Decoder Module: The decoder reconstructs the original multi-omics input data from the discrete embeddings. It consists of two stages: the integration decoder reconstructs the integrated feature representation from the discrete embeddings. The omics-specific decoders transform the integrated representation back into the respective omics data types, ensuring that the reconstructed data aligns with the original input features. 4) Classifier: To enhance the model’s ability to distinguish between different CRE types, CREATE includes a classifier that enforces the separation of CREs into distinct vectors in the codebook. CREs of the same type are encouraged to map to similar vectors, while those of different types are spread out across different vectors. This helps in achieving accurate and interpretable classifications.

CREATE offers several key advantages compared to existing methods: 1) Comprehensive Data Integration: By incorporating multiple omics data types, CREATE captures a more complete picture of the genomic context and CRE functionality. 2) Dynamic Codebook: The updateable codebook allows for flexible and accurate representation of CREs, overcoming limitations of fixed latent spaces in traditional VAE models. 3) Interpretable Embeddings: The discrete embeddings and their organization in the codebook provide clear and interpretable insights into CRE specificity and classification.

Overall, CREATE represents a significant advancement in computational CRE identification. Its ability to integrate multi-omics data, produce discrete embeddings, and offer interpretable results makes it a powerful tool for understanding gene regulation and its implications in complex biological processes.

### Cis-regulatory elements identification with CREATE

We comprehensively evaluated the performance of CREATE in identifying cell-type-specific CREs, including silencers, enhancers, promoters, insulators, and background regions, on the K562 and HepG2 cell types (Methods). To assess CREATE’s effectiveness, we conducted 10-fold cross-validation experiments and compared its performance with four baseline methods, including DeepSEA^17^, DanQ^18^, ES-transition^19^ and DeepICSH^20^. The primary evaluation metrics were area under the Receiver Operating Characteristic Curve (auROC), the area under the Precision-Recall Curve (auPRC) and the F1-score (Methods).

CREATE significantly surpasses the baseline methods by achieving the best classification performance on both K562 and HepG2 cell types (one-sided paired Wilcoxon signed-rank tests *P*-values < 1e-3), whereas the performance of baseline methods fluctuates across different cross-validation experiments (Fig. 2a-b and Supplementary Fig. 2a-b). For the K562 cell type, CREATE achieves a 10-fold macro-averaged auROC of 0.964 ± 0.002 (mean ± s.d.), outperforming the second-best method, ES-transition (0.928 ± 0.002) (Fig. 2c). Similarly, CREATE acquires a 10-fold macro-averaged auPRC of 0.848 ± 0.004, reflecting a substantial improvement of 10.5% compared to the second-best method, DeepICSH (0.743 ± 0.003) (Fig. 2d). A comparable performance enhancement is observed for the HepG2 cell type, with CREATE overtaking the baseline methods by a noticeable margin (Supplementary Fig. 2c-d).

**Fig. 2.**
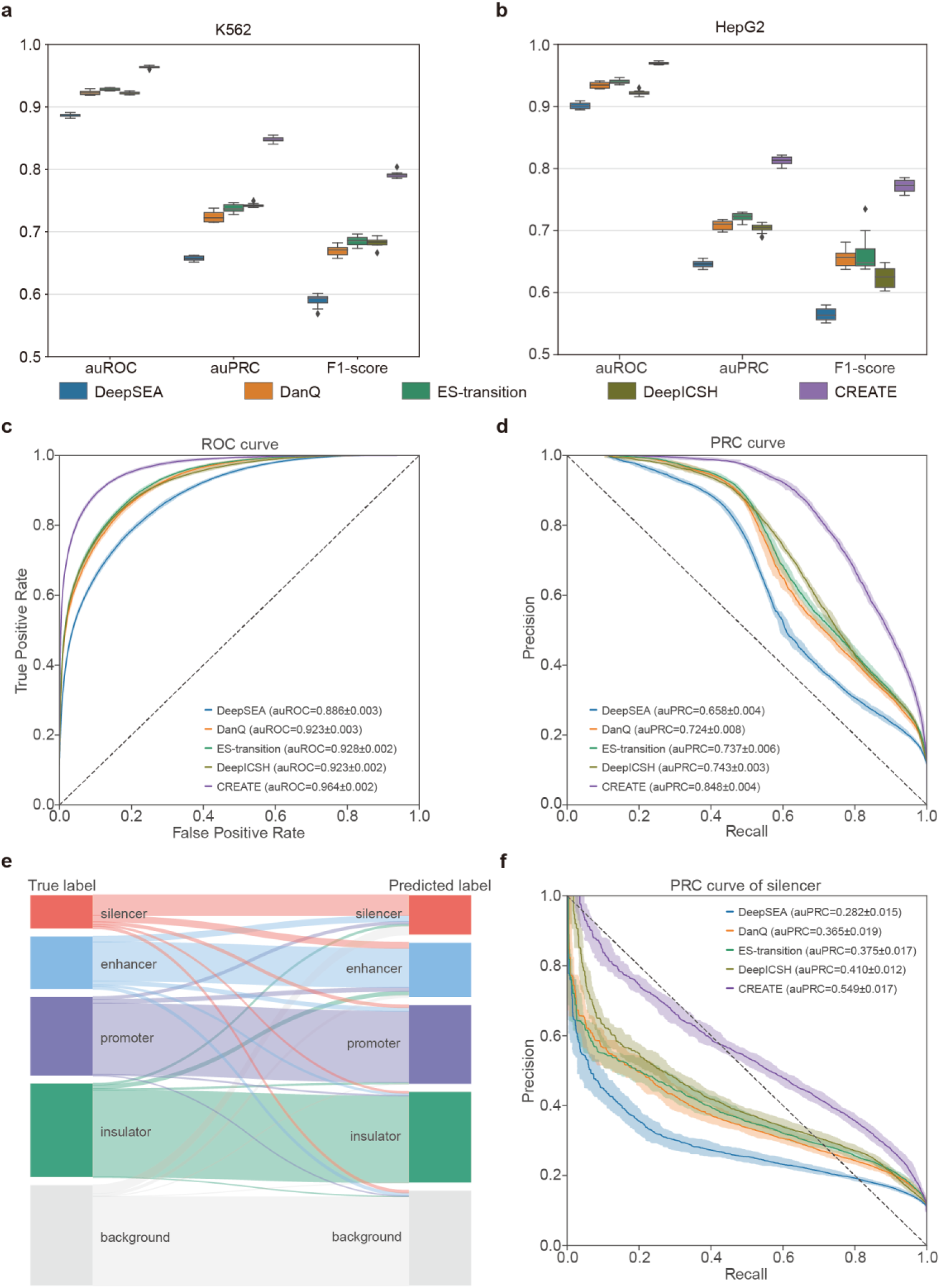
Evaluation of CREATE compared with the baseline methods. **a-b**, Boxplot of classification performance evaluated by auROC, auPRC and F1-score on the K562 cell type (**a**) and HepG2 cell type (**b**). Each box plot ranges from the upper to lower quartiles with the median as the horizontal line, whiskers extend to 1.5 times the interquartile range, and points represent outliers. **c**-**d**, Receiver Operating Characteristic curve (**c**) and Precision-Recall curve (**d**) comparing CREATE and baseline methods on the K562 cell type. **e**, The mapping between the true CRE labels and CREATE-predicted CRE labels on the testing data in one of the 10-fold cross-validation experiments of K562 cell type. **f**, Precision-Recall curve comparing CREATE and baseline methods for silencers in the K562 cell type. The mean and standard error of auROC or auPRC are reported in the legend. The confidence band shows ±1 s.d. for the averaged curve.

Among the various CRE types, silencers and enhancers present unique challenges due to their similar epigenetic signatures^31^. Despite this, CREATE demonstrated a clear distinction between these difficult-to-differentiate elements. For the K562 cell type, CREATE achieves notable improvements in identifying silencers, with a mean auPRC that was 13.9% higher than the second-best method, which shows a greater advantage than the macro-averaged results of all CREs (Fig. 2f and Supplementary Fig. 2e). Similarly, CREATE attains a remarkable 22.1% improvement in mean auPRC for enhancers compared with the second-best method (Supplementary Fig. 3a-b). For other CRE types—promoters, insulators, and background regions—CREATE also demonstrates optimal classification performance, although baseline methods provide competitive results (Supplementary Fig. 3c-h). The performance trends for the HepG2 cell type mirrored those observed for the K562 cell type, further validating CREATE’s robustness across different cell types (Supplementary Fig. 4).

The results highlight CREATE’s exceptional capability in accurately identifying and distinguishing between various CRE types, particularly those less studied or less abundant, such as silencers and enhancers. CREATE’s superior performance in capturing CRE variability and cell type specificity underscores its potential as a powerful tool for advancing our understanding of gene regulation mechanisms.

### Robustness and effectiveness of CREATE

CREATE integrates genomic sequences, chromatin accessibility, and chromatin interactions to deliver a thorough characterization of gene regulatory processes. To assess the contributions of these different inputs, we conducted extensive ablation experiments. We referred to the models employing a single type of omics data as CREATE(seq), CREATE(open) and CREATE(loop), and those incorporating two different types of omics data as CREATE(seq+open), CREATE(seq+loop) and CREATE(open+loop), respectively. Among the seven models evaluated, CREATE consistently demonstrates the highest classification performance, confirming the importance of incorporating chromatin accessibility and chromatin interactions for superior CRE identification (Fig. 3a and Supplementary Fig. 5a-b). Specifically, CREATE shows substantial improvements in identifying challenging CRE types such as silencers and enhancers (Supplementary Fig. 5c-d). Notably, CREATE(seq), which relies solely on genomic sequences, achieves a 10-fold macro-averaged auPRC of 0.800 ± 0.004, surpassing baseline methods by 5.7% in mean auPRC (Supplementary Fig. 5b). This underscores CREATE’s robust performance even when using genomic sequences alone. Incorporating additional omics data, such as chromatin accessibility or chromatin interactions, further enhances performance, though the inclusion of only these inputs without genomic sequences results in relatively poorer outcomes (Fig. 3a and Supplementary Fig. 5a). This indicates that while chromatin accessibility and interactions are valuable, genomic sequences are indispensable for optimal CRE identification, particularly contributing to better identification of silencers and enhancers.

**Fig. 3.**
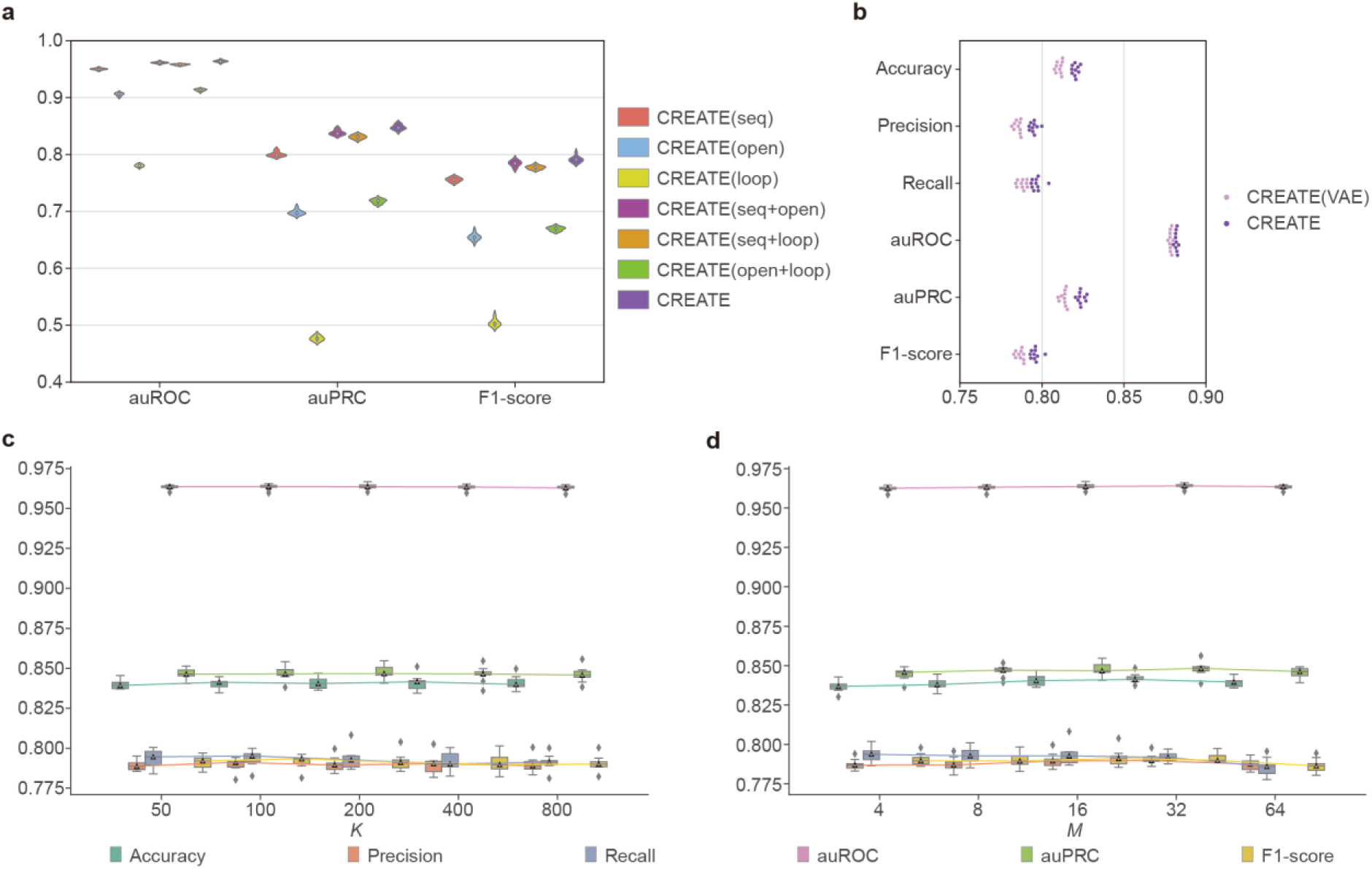
Robustness analysis of CREATE. **a**, Violin plot of classification performance evaluated by auROC, auPRC and F1-score for model ablation of CREATE on the K562 cell type. **b**, Swarm plot of classification performance evaluated by accuracy, precision, recall, auROC, auPRC and F1-score for CREATE compared with CREATE (VAE) on the K562 cell type. **c**, Classification performance of CREATE under different values of *K* (size of codebook) on the K562 cell type. **d**, Classification performance of CREATE under different values of *M* (time of split quantization) on the K562 cell type. Each box plot ranges from the upper to lower quartiles with the median as the horizontal line, whiskers extend to 1.5 times the interquartile range, and points represent outliers.

Based on the VQ-VAE framework^34, 35^, discrete embedding allows the latent space of CREATE to be a learnable discrete distribution, as opposed to the fixed Gaussian distribution as in VAE models^32^. To verify the efficiency of discrete embedding in CREATE, we compared CREATE with a variant using VAE latent space (CREATE(VAE)) while keeping other modules and training strategies unchanged. CREATE significantly outperformed CREATE(VAE) across all evaluation metrics (one-sided paired Wilcoxon signed-rank tests *P*-values < 1e-3) (Fig. 3b). This result highlights the effectiveness of discrete embeddings in capturing complex CRE features.

To validate the stability and effectiveness of CREATE, we designed comprehensive robustness analyses for the hyperparameters in CREATE, including *K* denoting the size of codebook, *M* denoting the time of split quantization^32, 36^, *α* denoting the weight of 𝐿_𝑒𝑛𝑐𝑜𝑑𝑒𝑟_, 𝜇 denoting the update ratio of codebook. First, to evaluate the robustness of CREATE to the size of codebook, we trained CREATE with different values of *K* (50, 100, 200, 400 and 800) on the K562 cell type. CREATE exhibited consistent classification performance across these values, demonstrating its insensitivity to codebook size variations (Fig. 3c). Taking into account the balance of CRE specificity preservation and codebook utilization, we set the default value of *K* to 200. Second, to evaluate the stability of CREATE to the time of split quantization, we trained CREATE with different values of *M* (4, 8, 16, 32 and 64) on the K562 cell type. The results show that CREATE attains highly stable classification performance across different values of *M* (Fig. 3d). Evidently, the lower the time of split quantization, the higher the dimension of codebook features. With the consideration that it is obviously challenging to look up the nearest neighbors for high-dimensional vectors, we set the default value of *M* to 16. Third, following the original studies of VQ-VAE, we aimed for the codebook to have less impact on the output of encoder so that we set the default value of *α*, the weight of 𝐿_𝑒𝑛𝑐𝑜𝑑𝑒𝑟_, to 0.25. To validate the robustness of CREATE with different weights of 𝐿_𝑒𝑛𝑐𝑜𝑑𝑒𝑟_, we trained CREATE with different values of *α* (0.05, 0.1, 0.25, 0.5 and 1.0) on the K562 cell type. The results demonstrate that CREATE consistently obtains stable classification performance under different values of *α* (Supplementary Fig. 5e). Fourth, similar to the original studies of VQ-VAE, we set the default value of 𝜇, the update ratio of codebook, to 0.01. To assess the stability of CREATE with different update ratios, we trained CREATE under a series of 𝜇, 0.001, 0.005, 0.01, 0.05 and 0.1, on the K562 cell type. The results demonstrate the stability of the classification performance under different values of 𝜇 (Supplementary Fig. 5f). To summarize, the effective integration of multiple omics inputs, stable hyperparameters, and efficient discrete embedding all demonstrate the robustness and usability of CREATE.

### Feature spectrum for unveiling CRE specificity

Discrete latent embedding of CREATE can reveal biological insights in an interpretable and intuitive manner. Using the latent embeddings of CREs, we built a uniform manifold approximation and projection (UMAP)^37^ plot (Fig. 4a). Clearly, promoters, insulators and background regions are effectively separated, while there is some degree of overlap between silencers and enhancers, which is consistent with the classification results. To further validate the capability of CREATE in quantitatively articulating CRE specificity, we obtained specific feature spectrum for each type of CRE (Supplementary Fig. 6a and Methods). Briefly, each element in the CRE-specific feature spectrum represents the probability of a codebook feature occurring in that particular CRE embeddings. We can always discover a set of particular features that are uniquely associated with a specific CRE and have the highest probability scores on that CRE, and we refer to these features as CRE-specific features. Concretely, for the K562 cell type, there are specific features uniquely enriched in the feature spectrum of each CRE (Fig. 4b). For example, we definitely observe different sets of specific features corresponding to promoters, insulators and background regions, which are clearly separated in the UMAP visualization (Fig. 4a) and Sankey diagram (Fig. 2e) as well. For the most difficult-to-distinguish two types of CREs, the feature spectrum of silencers contains a set of features (to the left of the blue dashed line) with notably higher probability scores compared to their scores in the feature spectrum of enhancers. Similarly, there is a set of features (between the blue dashed line and the purple dashed line) with notably higher probability scores in the feature spectrum of enhancers than those of silencers. In short, there is a relatively clear difference between the feature spectra of silencers and enhancers while they are connected together in the UMAP visualization. A similar result also occurred on the HepG2 cell type (Supplementary Fig. 6b-c). The CRE-specific feature spectrum, derived from discrete latent embedding of CREATE, has the potential to depict the general and comprehensive patterns of a type of CRE, further unveiling the CRE specificity quantitatively and interpretably.

**Fig. 4.**
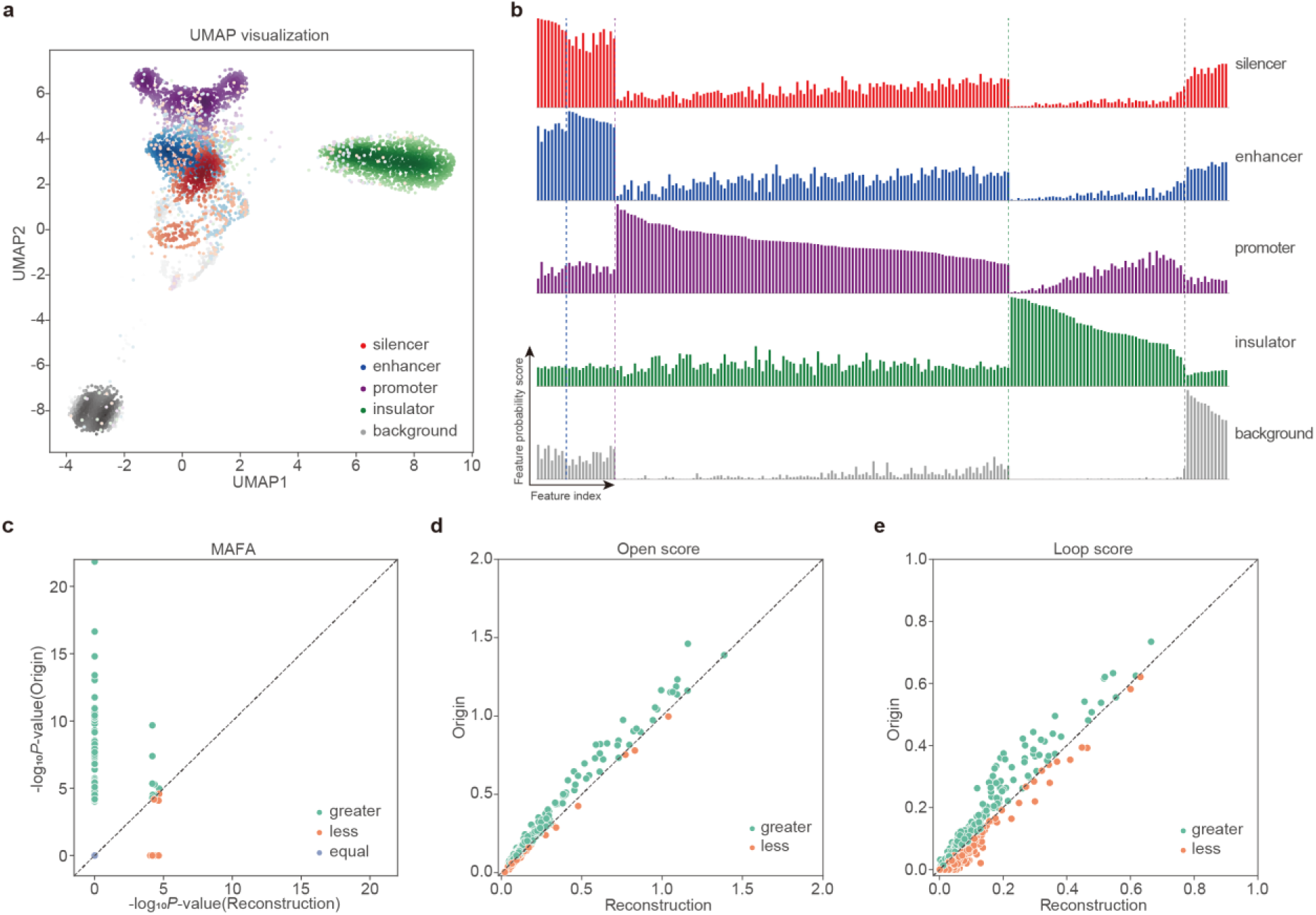
Generation and interpretation of CRE-specific feature spectrum. **a**, UMAP visualization of the CRE embeddings from CREATE on the testing data in one of the 10-fold cross-validation experiments of K562 cell type. **b**, CRE-specific feature spectrum. There are a distinct set of specific features that are enriched or depleted in the feature spectrum of each CRE on the K562 cell type. **c**, Comparison of MAFA motif enrichment significance (-log*_10_P*-value) between original input and reconstructed output when information derived from the major feature in the silencer-specific feature spectrum of K562 cell type is removed by zeroing it out before passing the CRE embeddings again through the decoder. **d**, Comparison of open scores between original input and reconstructed output when information derived from the major feature in the silencer-specific feature spectrum of K562 cell type is removed. **e**, Comparison of loop scores between original input and reconstructed output when information derived from the major feature in the silencer-specific feature spectrum of K562 cell type is removed.

To demonstrate the potential of codebook features in the CRE-specific feature spectrum for capturing key biological patterns, we identified the codebook feature with the highest probability score in the CRE-specific feature spectrum of K562 cell type as the major feature of that CRE, and we then zeroed it out before passing the CRE embeddings through the decoder again to generate the reconstructed output. To better understand the relationship between multi-omics input and the major feature of silencers, we designed comparative experiments between the original and reconstructed genomic sequences, chromatin accessibility scores and chromatin interaction scores. First, we conducted motif enrichment analysis for the original and reconstructed silencers (Methods). It is worth noting that the motif enrichment significance (-log_10_*P*-value) of MAFA, LHX6 and PAX8, which were reported as repressors in the previous literature^38–40^, is obviously higher in the original sequences compared to the reconstructed sequences (one-sided Wilcoxon rank-sum tests *P*-values < 7e-53) (Fig. 4c and Supplementary Fig. 6d-e), whereas similar comparison results were not observed for known activators, such as POU6F1^41^ and MYC^42^ (Supplementary Fig. 6f-g). This also demonstrates that the identified major feature of silencer-specific feature spectrum plays a crucial role in distinguishing between silencers and enhancers, as it indeed captures the motif information of some repressors, aligning with the repressive function of silencers. Simultaneously, TFs with the most significant difference between the original and reconstructed sequences, such as PRDM4, ZNF582 and SCRT2 (one-sided Wilcoxon rank-sum tests *P*-values < 6e-79) (Supplementary Fig. 7a-c), are considered to be novel silencer-related TFs. PRDM4 has been linked with recruiting chromatin modifiers, suggesting its involvement in establishing repressive chromatin states^43^. ZNF582 has been implicated in DNA methylation processes, which are crucial for maintaining silencer function^44^. SCRT2 is less characterized, but its differential binding indicates a possible regulatory role in silencing mechanisms^45^. Similarly, the CRE-specific motif information is also harbored in the major feature of enhancers, promoters and insulators (Supplementary Fig. 7d-i), demonstrating that these features catch CRE-specific sequence patterns. Additionally, the unique motif patterns associated with silencers compared to enhancers, promoters, and insulators provide further evidence that these elements are distinct regulatory modules with specific TF associations. This distinction underscores the importance of considering a broader scope of CREs, including dual-function regulatory elements that might act as silencers under certain conditions and enhancers under others. Furthermore, the zeroing operation led to a reduction in the reconstructed chromatin accessibility scores and chromatin interaction scores (Fig. 4d-e), indicating that the major feature also captures silencer-specific epigenomic characteristics. Through the extensive comparative experiments that we designed, the CRE-specific feature spectrum generated by CREATE interpretably reveals the CRE specificity and is potentially involved in the gene regulation process in specific cell types. In conclusion, CREATE not only identifies known regulatory elements but also sheds light on less understood elements like silencers, filling a critical gap in the current landscape of gene regulation studies.

### Large-scale prediction of cis-regulatory elements

The emergence of extensive epigenomic sequencing data across various cell types has enabled us to leverage a wealth of information for identifying cell-type-specific CREs on a large scale and establishing regulatory elements maps. CREATE proves to be a powerful tool for a comprehensive characterization of gene regulatory processes, revealing CREs with high accuracy and interpretability. In our study, we collected 270,259 candidate CREs on the K562 cell type and 232,456 candidate CREs on the HepG2 cell type for large-scale prediction (Supplementary Table 1 and Methods). For each cross-validation experiment, based on the trained CREATE model, we calculated a cutoff score for each type of CRE according to the validation set with a false positive rate (FPR) not exceeding 0.01. Candidate regions in Supplementary Table 1 exceeding the silencer cutoff score are marked as predicted silencers, and other CREs are labeled similarly. This approach led to the identification of 26,012 predicted silencers, 29,423 predicted enhancers, 2,057 predicted promoters, and 10,558 predicted insulators in the K562 cell type using the models trained on the K562 cell type, and the remaining sequences were classified as background regions. Similarly, in the HepG2 cell type, we identified 16,000 predicted silencers, 49,145 predicted enhancers, 4,422 predicted promoters and 13,122 predicted insulators using the models trained on the HepG2 cell type. The predicted CREs showed strong correlations with known epigenomic markers. For example, H3K27me3, a key histone modification associated with silencers^46–48^, was found at higher proportions in predicted silencers (9.0% in K562 and 16.0% in HepG2) compared to other CREs (average 2.2% in K562 and average 4.1% in HepG2) (highlighted with red dashed box; Fig. 5a and Supplementary Fig. 8a). Similarly, histone modifications associated with enhancers, such as H3K9ac^49, 50^, H3K27ac^50–52^, H3K4me1^50, 53, 54^, H3K4me2^54^ and H3K4me3^54, 55^, were more prevalent in predicted enhancers compared to other CRE types (average 18.5% in K562 and average 36.6% in HepG2) (highlighted with blue dashed box; Fig. 5a and Supplementary Fig. 8a).

**Fig. 5.**
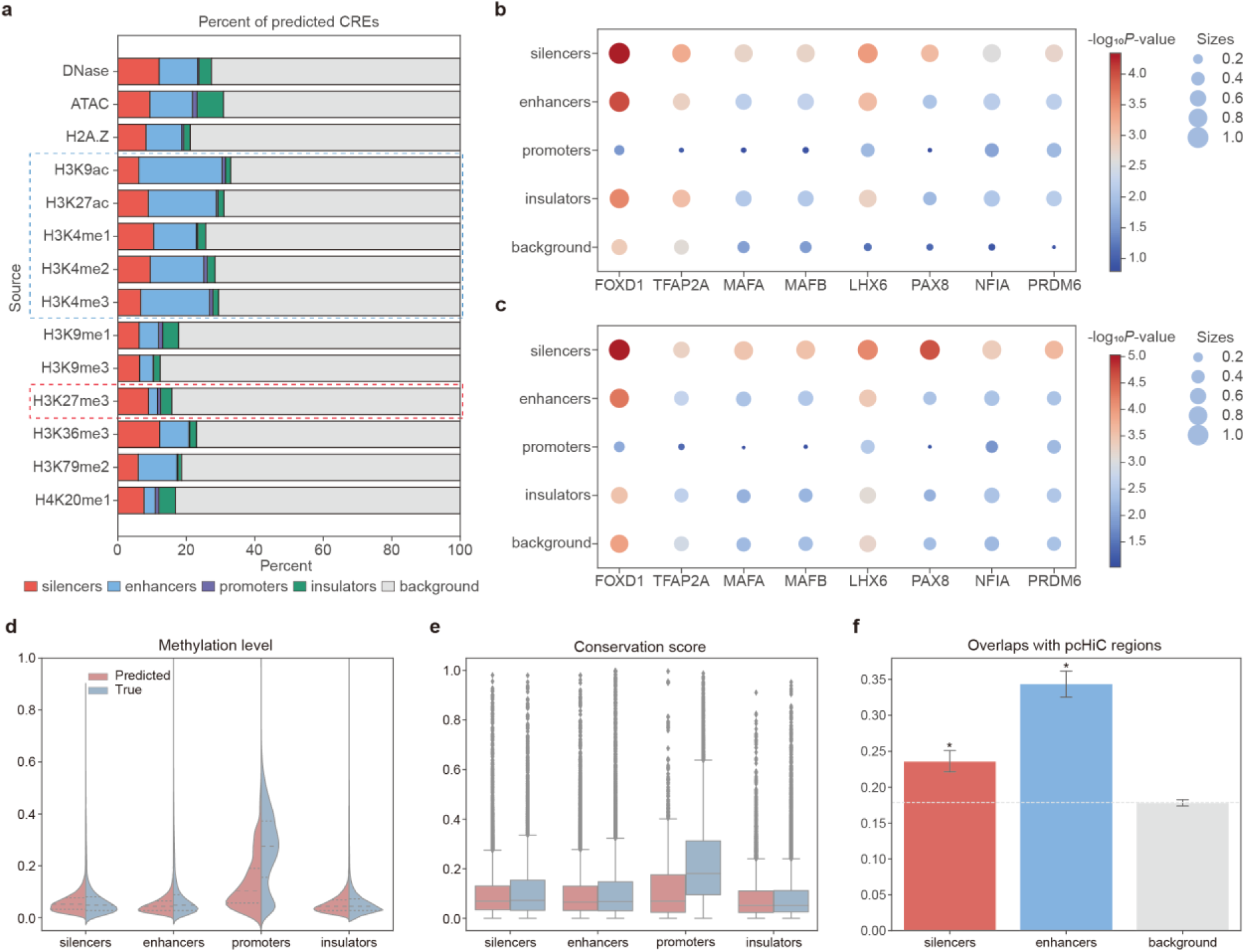
Characteristics of predicted CREs by CREATE. **a**, Percentage of predicted CREs and background regions from different candidate sources in the K562 cell type. **b-c**, Bubble plot of motif enrichment significance (-log*_10_P*-value) of repressive TFs (silencer-related TFs) at true CREs (**b**) and predicted CREs (**c**) on the K562 cell type. The size of bubbles represents the proportion of CREs with *P*-value < 0.01. **d**, Violin plot of methylation levels at true CREs and predicted CREs on the K562 cell type. Each violin plot contains three horizontal dashed lines denoting the median, the upper quartile, and the lower quartile. **e**, Box plot of conservation scores at true CREs and predicted CREs on the K562 cell type. Each box plot ranges from the upper to lower quartiles with the median as the horizontal line, whiskers extend to 1.5 times the interquartile range, and points represent outliers. **f**, Bar plot of overlaps between the pcHiC regions and predicted silencers, enhancers or background regions on the K562 cell type. The asterisks above the bars indicate the significant enrichments compared with the background regions. (∗) *P*-value < 2e-6. The error bars denote the 95% confidence interval, and the centers of error bars denote the average value.

To further validate the epigenomic characteristics of predicted CREs on a large scale, we conducted extensive comparative analyses. First, TFs play a crucial regulatory role in gene transcription by binding to CREs, and sequence-specific TF motifs can be considered key factors in identifying CREs^56–58^. We performed motif enrichment analysis on both true CREs (experimentally validated CREs) and predicted CREs (Methods). Compared to other true CREs and background regions, true silencers are enriched with the binding motifs of repressive TFs previously reported in the literature (Fig. 5b), such as FOXD1^59^, TFAP2A^60^, MAFA^38^, MAFB^61^, LHX6^39^, PAX8^40^, NFIA^62^ and PRDM6^63^, which are also enriched in the predicted silencers (Fig. 5c). Motifs belonging to active TFs, including POU6F1^41^, MYC^42^, ZFHX3^64^ and SOX8^65^, are enriched consistently across the true and predicted enhancers (Supplementary Fig. 8b-c). Notably, silencers and enhancers, the two most similar types of CREs, are enriched with the same TFs, such as MYC and ZFHX3. These TFs have been validated to act as either activators or repressors^42, 64, 66, 67^, which aligns with the potential conversion between silencers and enhancers under different conditions^19, 68^. Similar motif enrichment results, consistent between true and predicted CREs, have also been observed in promoters (Supplementary Fig. 8d-e). These results indicate that CREATE effectively captured the CRE-specific sequence characteristics.

Second, DNA methylation is an important epigenetic modification involved in gene regulation, particularly gene silencing^69, 70^. We calculated the methylation levels for both true CREs and predicted CREs (Methods), and observed the consistency between them except for promoters (Fig. 5d), which may be due to the complete collection of experimentally validated promoters and the limited number of predicted promoters. Specifically, 65.1% of the predicted promoters are adjacent to the promoters of non-coding genes, which is much higher than the 9.6% of randomly sampled genomic regions. In addition, the methylation levels of predicted CREs are significantly higher than those of predicted background regions (one-sided Wilcoxon rank-sum tests *P*-values < 2e-6) (Supplementary Fig. 8f).

Third, CREs are usually conserved in the evolutionary process of vertebrates, and the conserved regions are essential for deciphering the landscapes of gene regulation^71–73^. We computed the phastCons scores for the true CREs and the predicted CREs (Methods), and noticed that the conservation scores exhibit strong consistency between them except for promoters (Fig. 5e). Compared with the predicted background regions, the predicted CREs except insulators are significantly more conserved (one-sided Wilcoxon rank-sum tests *P*-values < 1e-13) (Supplementary Fig. 8g).

Fourth, CREs frequently regulate gene expression by connecting promoters through chromatin loops, which can be identified by promoter-capture HiC (pcHiC)^74^. We counted the number of overlaps between the pcHiC regions and the true CREs or the predicted CREs (Methods), and perceived the predicted silencers and enhancers harbor significantly more overlaps with pcHiC regions than the predicted background regions (one-sided Wilcoxon rank-sum tests *P*-values < 2e-6) (Fig. 5f), which aligns with the functional roles of these CREs in influencing target genes.

We further tested CREATE’s cross-cell type prediction capabilities by using models trained on the K562 cell type to predict CREs in the HepG2 cell type. The results (auROC of 0.964 ± 0.002 and auPRC of 0.792 ± 0.005) demonstrate that CREATE maintains high performance across different cell types, confirming its robustness and generalizability (Supplementary Fig. 2c-d). Collectively, CREATE precisely extracts the CRE-specific epigenomic characteristics, enabling the construction of a comprehensive CRE atlas.

### Characterization of dual-function regulatory elements

Dual-function regulatory elements (DFREs) are regions that exhibit dual roles as either silencers or enhancers depending on the cellular context^9, 19, 68, 75^. Understanding DFREs is crucial for unraveling the complexity of gene regulation, as these elements can significantly impact gene expression by switching functions based on the cellular environment. A K562 silencer overlapping a HepG2 enhancer by more than 600 bp was considered a DFRE, resulting in 2,409 DFREs (9.3% of all predicted silencers) and 23,603 normal silencers. Conversely, we identified 36,448 HepG2 enhancers that do not function as enhancers or silencers in the K562 cell type, categorizing them as normal enhancers. Reasonably, the ability of CREATE to effectively characterize DFREs is demonstrated by its capacity to assign higher CREATE enhancer scores to DFREs than normal silencers (one-sided Wilcoxon rank-sum test *P*-value < 6e-47) (Fig. 6a), and higher CREATE silencer scores than normal enhancers (one-sided Wilcoxon rank-sum test *P*-value < 2e-5) (Fig. 6b). This differentiation underscores CREATE’s effectiveness in distinguishing between multifunctional and context-specific regulatory elements.

**Fig. 6.**
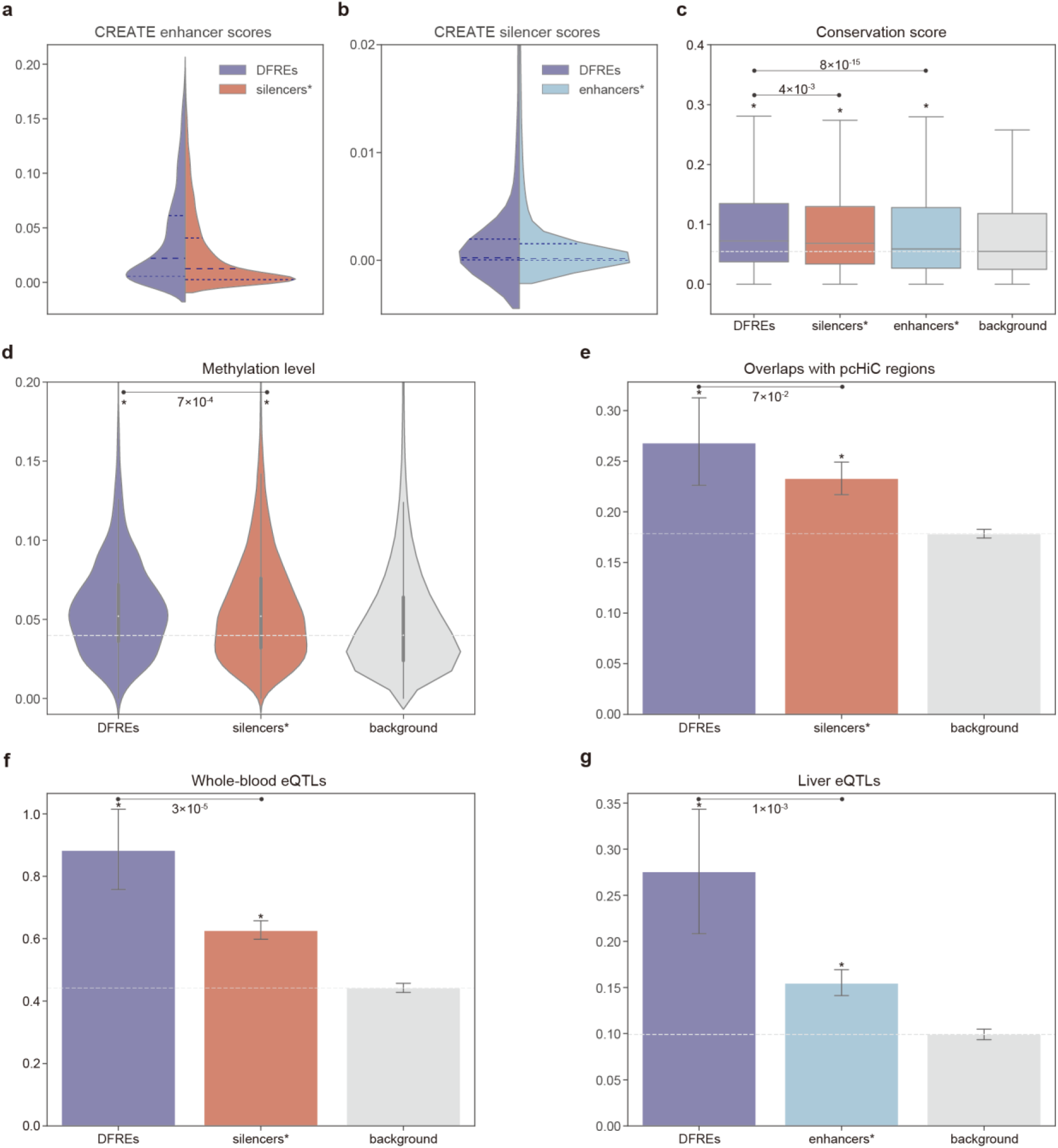
Characterization of DFREs identified by CREATE. **a**, Violin plot of the CREATE enhancer scores for DFREs and normal silencers (silencers*). **b**, Violin plot of the CREATE silencer scores for DFREs and normal enhancers (enhancers*). Each violin plot contains three horizontal dashed lines denoting the median, the upper quartile, and the lower quartile. **c**, Box plot of conservation scores at DFREs, normal silencers (silencers*), normal enhancers (enhancers*) and the background regions. The asterisks above the boxes indicate the significant enrichments compared with the background regions. (∗) *P*-value < 2e-31. Each box plot ranges from the upper to lower quartiles with the median as the horizontal line, whiskers extend to 1.5 times the interquartile range. **d**, Violin plot of methylation levels at DFREs, normal silencers (silencers*) and the background regions. (∗) *P*-value < 2e-6. **e**, Bar plot of overlaps between the pcHiC regions and DFREs, normal silencers (silencers*) or the background regions. (∗) *P*-value < 2e-3. **f**, Bar plot of overlaps between the whole-blood eQTLs and DFREs, normal silencers (silencers*) or the background regions. (∗) *P*-value < 5e-21. **g**, Bar plot of overlaps between the liver eQTLs and DFREs, normal enhancers (enhancers*) and the background regions. (∗) *P*-value < 2e-6. The error bars denote the 95% confidence interval, and the centers of error bars denote the average value.

To further explore the biological significance of DFREs, we conducted several comparative analyses. First, DFREs exhibit the highest conservation scores (one-sided Wilcoxon rank-sum tests *P*-values < 4e-3) and the background regions have the lowest conservation scores (one-sided Wilcoxon rank-sum tests *P*-values < 2e-31) (Fig. 6c), suggesting that these elements are evolutionarily preserved due to their critical roles in gene regulation. This high conservation underscores their functional importance across species and reinforces the value of identifying these elements for understanding gene regulatory mechanisms. Second, DFREs possess higher methylation levels in the K562 cell type compared to normal silencers (one-sided Wilcoxon rank-sum test *P*-value < 7e-4) (Fig. 6d). This observation highlights the unique epigenomic signatures of DFREs, suggesting that their dual functionality is associated with distinct methylation patterns, which may influence their regulatory roles. Third, DFREs show a strong preference with more overlaps with pcHiC regions of K562 cell type than normal silencers (Fig. 6e). This indicates that DFREs are actively involved in chromatin looping interactions, which are critical for mediating gene expression and regulatory network organization. Fourth, we computed the number of overlaps with expression quantitative trait loci (eQTLs) from whole-blood or liver tissues from GTEx^76, 77^ for DFREs, normal silencers, normal enhancers and the background regions (Methods). DFREs are significantly enriched with more whole-blood eQTLs than normal silencers (one-sided Wilcoxon rank-sum test *P*-value < 3e-5) (Fig. 6f), and more liver eQTLs than normal enhancers (one-sided Wilcoxon rank-sum test *P*-value < 2e-3) (Fig. 6g). This enrichment demonstrates the tissue-specific regulatory potential of DFREs, highlighting their role in fine-tuning gene expression across different biological contexts. Ultimately, CREATE’s advanced capability to differentiate and interpret DFREs enriches the field’s ability to map regulatory landscapes and uncover the underlying mechanisms of gene regulation.

### Disease-associated variations analysis and tissue-specific enrichments in CREs

CREs play crucial roles in disease susceptibility and phenotype variations, often harboring single-nucleotide polymorphisms (SNPs) and eQTLs associated with various diseases and traits^78, 79^. To highlight the capacity of CREATE in uncovering disease-relevant variations within CREs, we analyzed overlaps with SNPs from dbSNP^80–82^ database and eQTLs from GTEx^76, 77^ for both the true CREs and the predicted CREs (Methods). Our results reveal that the gene variation distributions at predicted CREs align closely with those at true CREs (Fig. 7a and Supplementary Fig. 9a-c). In detail, the predicted CREs are significantly enriched with more rare SNPs than the background regions (one-sided Wilcoxon rank-sum tests *P*-values < 2e-9) (Fig. 7b), whereas a similar significant result was not observed for common SNPs (Supplementary Fig. 9d), suggesting the functional importance of identified CREs. The enrichment of rare gene variants in CREs, especially silencers and enhancers, supports the notions that disease-associated variants are more frequently located in gene regulatory regions^9, 12, 83^, and rare variants are more impactful in complex diseases compared to common gene variants^84, 85^. Similarly, compared to background regions, significant enrichment on silencers and enhancers also occurred with whole-blood eQTLs (one-sided Wilcoxon rank-sum tests *P*-values < 4e-49) (Supplementary Fig. 9e), but not with all eQTLs (Supplementary Fig. 9f), reinforcing their tissue specificity. Besides, gene variation levels for both true CREs and predicted CREs gradually decrease with an increase in CREATE background scores (Fig. 7c and Supplementary Fig. 10a,d-i) but increase with an increase in CREATE silencer scores (Supplementary Fig. 10b-c), explicating the ability of CREATE for quantifying the impact of gene variations.

**Fig. 7.**
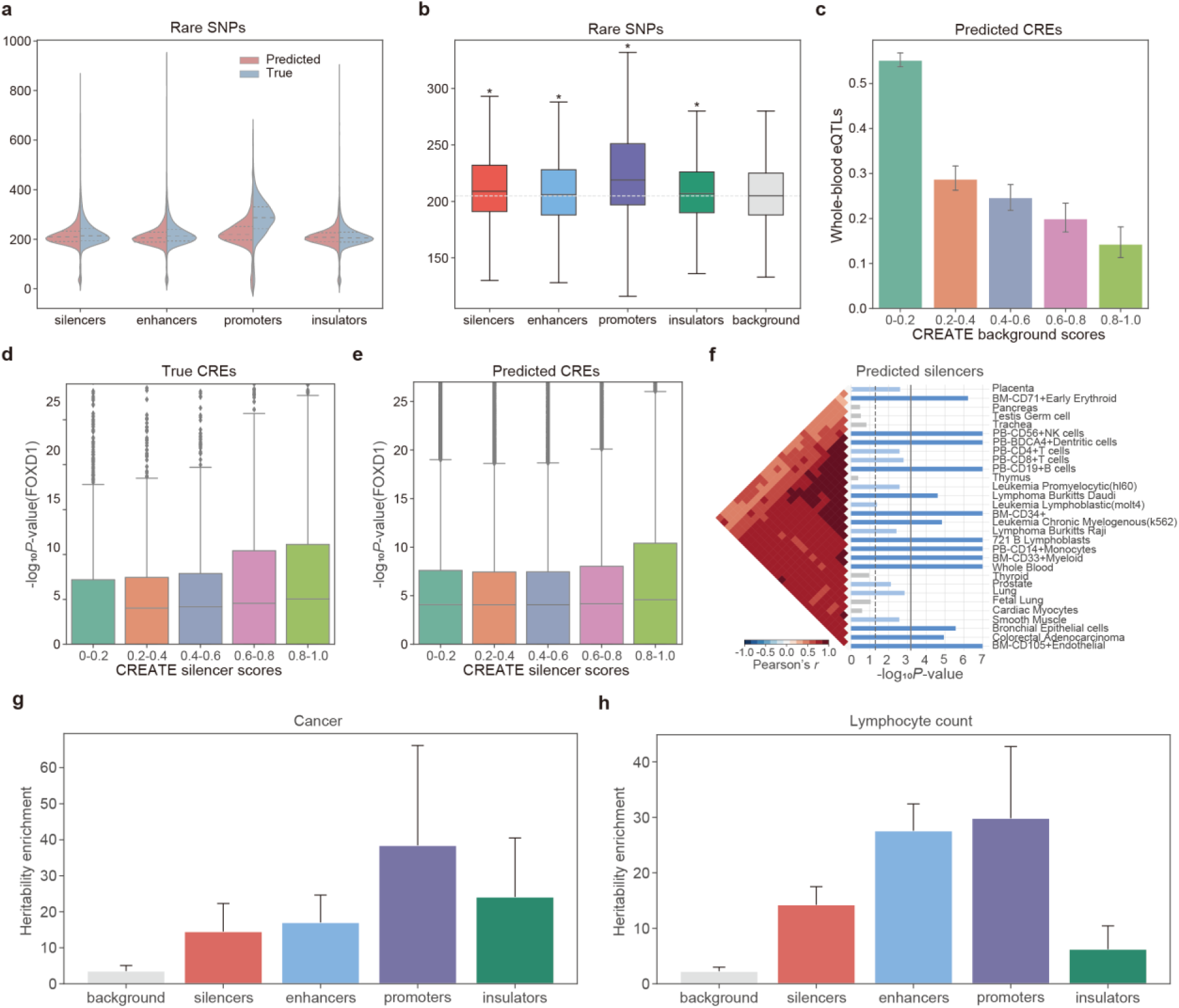
Identification of the biological variability of CREs by CREATE. **a**, Violin plot of overlaps between the rare SNPs and true CREs or predicted CREs on the K562 cell type. Each violin plot contains three horizontal dashed lines denoting the median, the upper quartile, and the lower quartile. **b**, Box plot of overlaps between the rare SNPs and the predicted CREs or background regions on the K562 cell type. The asterisks above the boxes indicate the significant enrichments compared with the background regions. (∗) *P*-value < 2e-9. Each box plot ranges from the upper to lower quartiles with the median as the horizontal line, whiskers extend to 1.5 times the interquartile range. **c**, Correlation between the CREATE background scores and overlaps between the whole-blood eQTLs and predicted CREs on the K562 cell type. **d-e**, Correlation between the CREATE silencer scores and the motif enrichment significance (-log*_10_P*-value) of FOXD1 at true CREs (**d**) and predicted CREs (**e**) on the K562 cell type. Each box plot ranges from the upper to lower quartiles with the median as the horizontal line, whiskers extend to 1.5 times the interquartile range, and points represent outliers. **f**, Top 30 significantly enriched tissues in SNPsea analysis on the predicted silencers of K562 cell type. The vertical dashed line represents the one-sided *P*-value cutoff at the 0.05 level, while the solid lines denotes the cutoff at 0.05 level for the one-sided *P*-value with Bonferroni correction. Each plot also contains the ordered expression profiles using hierarchical clustering with unweighted pair-group method with arithmetic means, and the Pearson correlation coefficients indicating the correlation between profiles. **g-h**, Heritability enrichments estimated by LDSC within predicted CREs and background regions identified by CREATE for blood-related traits including cancer (**g**) and lymphocyte count (**h**). The error bars denote jackknife standard errors over 200 equally sized blocks of adjacent SNPs about the estimates of enrichment, and the centers of error bars represent the average value.

To further quantitatively assess the power of CREATE in unlocking the CRE-specific sequence characteristics, we portrayed the correlation between the CREATE scores and the motif enrichment significance at true or predicted CREs (Fig. 7d-e and Supplementary Figs. 11-14). We recognized the strong positive correlation between the CREATE silencer scores and the enrichment significance of silencer-related TFs (Fig. 5b-c), as well as between the CREATE enhancer scores and the enrichment significance of enhancer-related TFs (Supplementary Fig. 8b-c).

To illustrate the ability of CREATE for revealing the tissues influenced by the identified risk loci within the true CREs and the predicted CREs, we applied SNPsea^86^ for tissue enrichment analysis (Methods). For the silencers and enhancers predicted by CREATE, we discovered more tissues related to blood than background regions (Fig. 7f and Supplementary Fig. 15), which aligns with the outcomes of true CREs (Supplementary Fig. 16). Concretely, Leukemia_Chronic_Myelogenous(k562) was identified as a significantly enriched tissue for the predicted silencers and enhancers in the K562 cell type (*P*-values < 2e-5), confirming the tissue specificity of CREs identified by CREATE.

To validate the competence of CREATE for studying the variations in phenotypes based on the true CREs and the predicted CREs, we utilized partitioned linkage disequilibrium score regression (LDSC)^87^ for heritability enrichment analysis (Methods). Specifically, the enrichment of heritability in the predicted CREs is higher than that in the predicted background regions for the blood-related phenotypes, such as cancer, lymphocyte count and so on (Fig. 7g-h and Supplementary Fig. 17). Along with the enrichment results for the true CREs and background regions (Supplementary Fig. 18), CREs predicted by CREATE have inherited the pattern of heritability contribution for complex traits and diseases.

Altogether, CREATE excels in identifying CREs with significant disease-related variations and tissue-specific enrichments, providing critical insights into regulatory dynamics during development and disease progression. This capability underscores the power of CREATE in advancing our understanding of gene regulation and its implications for complex traits and diseases.

## Discussion

In this study, we introduce CREATE, a groundbreaking multimodal architecture that integrates DNA sequences, cell-type-specific chromatin accessibility, and chromatin interaction features for the multi-class prediction of CREs. Utilizing discrete CRE embeddings, we have verified the superior performance of CREATE in accurate CRE identification compared to state-of-the-art methods, as well as its effectiveness and stability across various input combinations, hyperparameters and forms of latent space. One of the key strengths of CREATE lies in its ability to offer improved interpretability as the CRE-specific feature spectrum, which quantitatively elucidates the CRE specificity and captures CRE-specific epigenomic characteristics. Moreover, CREATE has been validated the substantial potential in identifying cell-type-specific CREs on a large scale and uncovering biological variabilities of CREs, illustrating the ability of CREATE for unveiling the underlying regulatory dynamics that drive transcriptional regulation and disease development.

However, despite its successes, there are several areas where CREATE could be further improved. Our study identifies a few limitations and suggests several future directions for enhancing the method: 1) Data imbalance and insufficient research on certain CREs. The current data imbalance, particularly for silencers and certain cell types, impairs the overall performance of CRE identification. The obscure understanding of general silencer characteristics, the limited number of experimentally validated silencers and the restricted number of cell types studied, pose challenges in both model training and the selection of background regions. To address this, we plan to update our predicted silencers to the SilencerDB database^88^ and expand our identification to include more cell types. We also anticipate that advances in biological technologies, such as HiChIP with a broader range of ChIPs, will enhance multi-class CRE identification and aid in constructing a more comprehensive regulatory atlas. 2) Per-base-paired input features and input-specific encoder-decoder structure. While the per-base-paired input features and the input-specific encoder-decoder structure are effective for extracting detailed and comprehensive CRE embeddings, some epigenetic features, such as TF binding, are not well-represented in this format. To improve scalability and representation, we propose integrating prior biological knowledge as additional constraints directly applied to the codebook. This approach is expected to enhance the model’s ability of capturing complex epigenetic features and improve overall performance. 3) Development of a unified foundation model. Gene regulatory analysis methods typically focus on specialized models for specific problems. This paradigm limits the generalizability and integration of findings across different contexts. We aspire to develop a foundation model for the unified characterization of key gene regulatory factors, leveraging the shareability, scalability and interpretability of the discrete embedding in CREATE. We anticipate that such a foundation model will facilitate a deeper understanding of gene regulation mechanisms and their implications for disease development, ultimately enabling biological discoveries and applications in developmental biology and precision medicine.

In conclusion, CREATE represents a significant advancement in the prediction and interpretation of CREs, offering superior performance and insights compared to existing methods. Its ability to integrate diverse data types and deliver interpretable results positions it as a valuable tool for exploring gene regulation and disease mechanisms. Future improvements and expansions of CREATE will continue to refine its capabilities and extend its applicability, driving forward our understanding of the complex interplay between gene regulation and disease.

## Methods

### Data collection and preprocessing

All datasets used in this study were publicly available and collected from different sources. We downloaded experimentally validated silencers for K562 and HepG2 cell types from the SilencerDB database^88^. We downloaded experimentally validated enhancers for K562 and HepG2 cell types from the FANTOM5 project^24, 26^. We obtained transcription start sites (TSSs) from the EPD database^89^ and defined 1kb regions surrounding TSSs (500 bp upstream and 500 bp downstream) as promoters. Since CTCF characterized as an insulator by blocking chromatin interactions^90, 91^, we took as insulators the CTCF Chromatin immunoprecipitation sequencing (ChIP-seq) peaks for K562 and HepG2 cell types collected from the ENCODE project^92, 93^.

In addition, we collected multiple histone modification ChIP-seq peaks and chromatin accessibility peaks for K562 and HepG2 cell types from the Roadmap project^94^ and ENCODE project (Supplementary Table 1). After filtering the regions overlapped with the experimentally validated CREs, known genes and consensus black list, we obtained 270,259 and 232,456 candidate CREs for large-scale prediction on the K562 and HepG2 cell types, respectively.

We randomly sampled DNA sequences from the entire human reference genome, excluding the experimentally validated and candidate CREs, known genes, consensus black list. After filtering overlapping regions between CREs, we obtained 6754 silencers, 10,528 enhancers, 15,699 promoters, 18,631 insulators and 20,000 background regions for the K562 cell type, and 1456 silencers, 11,407 enhancers, 14,535 promoters, 15,650 insulators and 20,000 background regions for the HepG2 cell type. The input for each CRE comprises three components: a one-hot encoded 1000-bp sequence from the human GRCh37/hg19 reference genome, a vector containing chromatin open scores per base pair, and another vector containing chromatin loop scores per base pair.

### Chromatin open score

Chromatin accessibility is pivotal for identifying CREs, given that active regulatory DNA elements are typically situated in accessible chromatin regions^5, 6^. To incorporate the information of chromatin accessibility, we adopted OpenAnnotate^95^ to efficiently calculate the raw read open scores of CREs and background regions per base pair. We derived the chromatin open score per base pair by averaging the raw read open scores across replicates for each respective cell type.

### Chromatin loop score

Chromatin looping interactions exert a substantial influence on gene regulation by establishing connections between regulatory elements and target genes^7, 8^. We incorporated cell-type-specific chromatin interaction data from HiChIP, which precisely profiles both regulatory and structural interactions^16, 96^, to enhance the identification of CREs. We first calculated the number of chromatin loops per base pair for each CRE, and then obtained the chromatin loop score after logarithmic transformation.

### Data augmentation

To ensure enough training samples for our model, we applied a data augmentation strategy to CREs^21, 22, 97^. As illustrated in Supplementary Fig. 1, for each CRE with length of 1000 base pairs, we shifted a window along the reference genome with a stride of 10 from the midpoint towards both ends. To mitigate the impact of data imbalance, we optionally incorporated data augmentation with varying augmentation ratios (5:5:3:3:1) for silencers, enhancers, promoters, insulators and background regions in the training data. Additionally, we augmented CREs by including the reverse complement of each original sequence. To prevent information leakage, the augmentation ratios for CREs in the validation and testing data are kept consistent at 5. Take the average of the predicted probabilities for all augmented sequences of the input sequence as the predicted probability for that input sequence.

### The CREATE framework

We fed CREATE with a concatenated vector 𝐗^𝑖^ ∈ ℝ^6×𝐿^ for the *i*-th input sample including a one-hot encoded genomic sequence 𝐒^𝑖^ ∈ ℝ^4×𝐿^, a chromatin open score vector 𝐎^𝑖^ ∈ ℝ^1×𝐿^ and a chromatin loop score vector 𝐋^𝑖^ ∈ ℝ^1×𝐿^, where *L* is the length of sequence (*L*=1000). CREATE comprises encoders, a vector quantization module, and decoders. The encoder module of CREATE includes encoders for multiple input-specific learning and an encoder for integrating multiple inputs. Each encoder consists of a convolutional layer, a max-pooling layer, a ReLU non-linear activation function and a dropout layer. Correspondingly, each decoder consists of a deconvolutional layer, an upsample layer and a Sigmoid or ReLU non-linear activation function. In addition, we introduced a classifier with three fully connected layers to predict CREs based on their embeddings. Specifically, the output of the encoder module is denoted as 𝐞^𝑖^ ∈ ℝ^𝐿′×𝐷′^ for the *i*-th CRE, where 𝐿^′^ and 𝐷^′^ are the length and dimensionality of the latent embedding respectively, and after split quantization^32, 36^, it will be split into 𝐿^′^ × 𝑀 vectors 𝐞^𝑖^ ∈ ℝ^𝐷^, 𝑙 ∈ {1, …, 𝐿^′^}, 𝑗 ∈ {1, …, 𝑀}, where 𝑀 is the time of split quantization. Utilizing a shared codebook 𝐯_𝑘_ ∈ ℝ^𝐷^, 𝑘 ∈ {1, …, 𝐾} with the size of *K*, we obtained the quantized latent embedding 𝐪^𝑖^ ∈ ℝ^𝐿′×𝐷′^ for the *i*-th CRE by substituting the vector 𝐞_*ij*_^𝑖^ with the nearest counterpart in the codebook as follows:

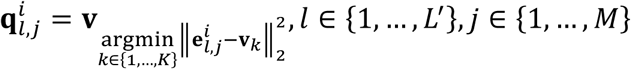

### Model training

We employed multiple update methods for different components of CREATE, mirroring the approach taken in the original studies of VQ-VAE^34, 35^. Let ℬ_0_ be a mini-batch of data for training.

First, to optimize the decoder and encoder by reducing the distance between the original input and the reconstructed output, we integrated a hybrid reconstruction loss comprising multiple components corresponding to different inputs:

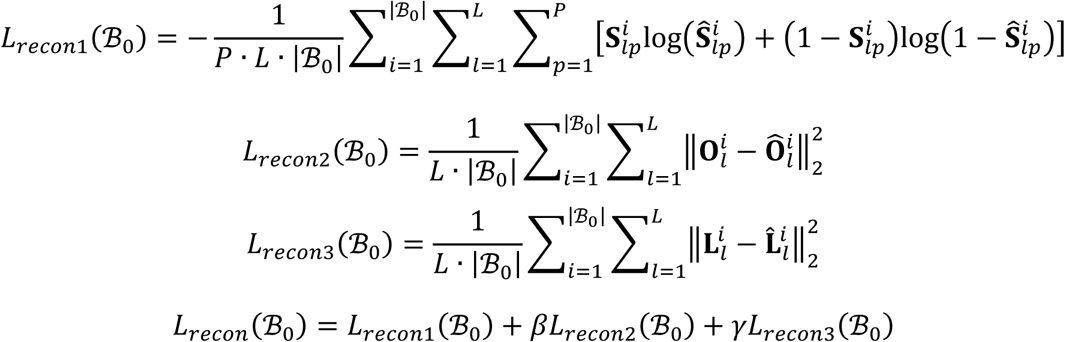

where *P* represents the number of different types of bases in the DNA sequence (*P*=4), 𝛽 and 𝛾 are the weights of 𝐿_𝑟𝑒𝑐𝑜𝑛2_ and 𝐿_𝑟𝑒𝑐𝑜𝑛3_ respectively (𝛽 = 0.01, 𝛾 = 0.1).

Second, to promote the encoder output to closely align with the selected codebook features and avoid excessive fluctuation, we introduced the encoder loss to aid in updating the encoder:

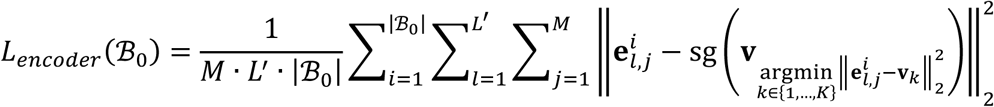

where sg(·) denotes the stop-gradient operator with zero partial derivatives.

Third, we followed the recommendation from both the original studies of VQ-VAE and recent related researches^98–101^ to utilize exponential moving average (EMA) for updating the codebook. Considering 𝑛_𝑘_ is the number of vectors matched to 𝐯_𝑘_ and 𝐞*_k_*_,𝑚_^∗^ is the *m*-th vector, we directly took the mean of the vectors in the set {𝐞*_k_*_,𝑚_^∗^ |𝑚 = 1, …, 𝑛_𝑘_} to optimize the code 𝐯_𝑘_ as follows:

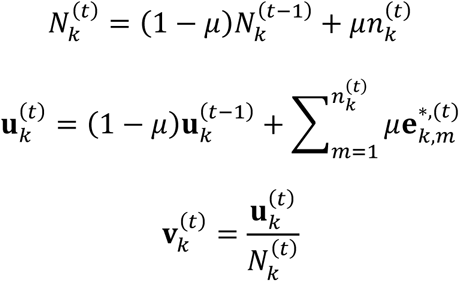

where 𝜇 is the update ratio of codebook. We initialized 𝑁_𝑘_ as a zero vector and 𝐮_𝑘_ randomly from a normal distribution with a mean of 0 and a standard deviation of 1.

Fourth, we further incorporated a classifier based on the CRE embeddings, with the cross-entropy loss function given by:

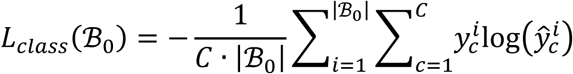

where *C* represents the number of types of CREs (*C*=5). To sum up, we trained CREATE using EMA and the total loss function as follows:

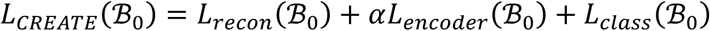

where *α* is the weights of 𝐿_𝑒𝑛𝑐𝑜𝑑𝑒𝑟_ .

In this study, we implemented CREATE with “Pytorch” package^102^. In details, there are three one-dimensional convolutional layers (filters=256,128,128; size=8,8,8) with layer normalization in the input-specific encoder module, followed by three one-dimensional convolutional layers (filters=512,384,128; size=1,8,8). In all cases, we set the mini-batch size to 1024 and employed the Adam stochastic optimization algorithm^103^ with a learning rate of 5e-5. We trained CREATE with a maximum of 300 epochs and implemented early stopping if there were no reductions in validation auPRC for 20 consecutive epochs. We set the dimension of the latent embedding to 128 and trained CREATE with *M* of 16, *K* of 200, *α* of 0.25, and *μ* of 0.01.

### Model evaluation

To comprehensively evaluate the performance of CREATE for CRE identification, we conducted 10-fold cross-validation experiments by dividing all CREs into 8:1:1 ratios for training, validation and testing data, respectively. We evenly distributed each type of CRE into 10 folds. We compared the classification performance with four baseline methods including DeepSEA^17^, DanQ^18^, ES-transition^19^ and DeepICSH^20^, with the area under the Receiver Operating Characteristic Curve (auROC), the area under the Precision-Recall Curve (auPRC), F1-score, accuracy, precision and recall as evaluation metrics.

### Feature spectrum

Supplementary Fig. 6a illustrates the process of generating the feature spectrum. For the *j*-th codebook feature, we counted its occurrence frequency in the latent embeddings of input regions, and we summed over these frequencies across all regions of the *i*-th CRE to gain the frequency 𝑐_𝑖𝑗_. We next derived a probability matrix (𝐏 ∈ ℝ^𝐶×𝐾^) by the following formula:

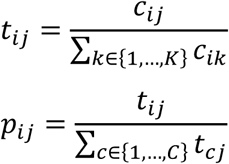

where *C* is the number of types of CREs and *K* is the number of codebook features. In this matrix, a row corresponds to a type of CRE and a column to a codebook feature, and an element 𝑝_𝑖𝑗_ indicates a feature probability score, representing the likelihood of the *j*-th codebook feature appearing in the latent embeddings of the *i*-th CRE. For the *j*-th codebook feature, we identified the element 𝑝_𝑗_ with the highest feature probability score and the corresponding CRE 𝐶_𝑗_ yielding the score, as follows.

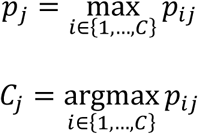

We then grouped the features corresponding to the same CRE together based on their CRE indices (*Cj*), and further sorted these features in descending order according to their feature probability scores (*p_j_*). Finally, we attained the rearranged matrix 𝐅 ∈ ℝ^𝐶×𝐾^ as the interpretable feature spectrum.

### Downstream analyses

#### Motif enrichment analysis

To discover enriched TF motifs for the true CREs and the predicted CREs by CREATE, we applied the tool FIMO^104^ with default settings to scan a set of input sequences for searching known human TFs in the HOCOMOCO^105^ database. For each input sequence, we used Fisher’s method to combine the *P*-values of reported binding sites for each TF, and we obtained a *P*-value vector representing the significance that 678 human TFs matched in the input sequence.

#### Methylation levels computation

The methylation state data at CpG in the K562 cell type was obtained from ENCODE^106^ (https://www.encodeproject.org/files/ENCFF867JRG/; https://www.encodeproject.org/files/ENCFF721JMB/). Using BEDTools^107^, we computed the methylation levels for the true CREs and the predicted CREs by CREATE.

#### Conservation scores computation

We downloaded the phastCons^71, 108^ scores for multiple alignments of 45 vertebrate genomes to the human genome from UCSC^109, 110^ (http://hgdownload.soe.ucsc.edu/goldenPath/hg19/phastCons46way/vertebrate.phastCons46way.bw). The phastCons scores for the true CREs and the predicted CREs were calculated via UCSC tool *bigWigAverageOverBed*^111^.

#### Overlaps with promoter-capture HiC regions

The pcHiC data of K562 cell type was downloaded from NCBI^112^ under accession number “GSE236305”. Using BEDTools^107^, we computed the number of overlaps between the pcHiC regions and the true CREs or the predicted CREs by CREATE.

#### Gene variation analysis

We downloaded human all SNPs and common SNPs from dbSNP^80–82^ database (https://ftp.ncbi.nih.gov/snp/organisms/human_9606_b151_GRCh37p13/VCF/), all GTEx eQTLs, whole-blood eQTLs and liver eQTLs from the Genotype-Tissue Expression Project^76, 77^, GTEx database version 7 (https://www.gtexportal.org/home/downloads/adult-gtex). By excluding the common SNPs from all human SNPs, we obtained rare SNPs. Using BEDTools^107^, we considered the number of overlaps with SNPs or eQTLs as the corresponding gene variation levels for the true CREs or the predicted CREs by CREATE.

#### Tissue enrichment analysis

To identify the tissues influenced by the identified risk loci within the true CREs and the predicted CREs by CREATE, we performed SNPsea analysis^86^ with default settings. Based on the tissue-specific expression profiles of 17,581 genes across 79 human tissues (Gene Atlas^113^), we quantified the enrichments of these profiles on the true CREs and the predicted CREs, and displayed the top 30 significantly enriched tissues in the heatmaps.

#### Heritability enrichment analysis

To quantify the enrichment of heritability for blood-related phenotypes within the true CREs and the predicted CREs by CREATE, we conducted heritability enrichment analysis using partitioned LDSC^87^ with default settings. LDSC took European samples from the 1000 Genomes Project as the LD reference panel. We downloaded the HapMap3 SNPs and GWAS summary statistics from the Broad LD Hub (https://doi.org/10.5281/zenodo.7768714), and then quantified the enrichment of heritability for blood-related phenotypes, and displayed the results for the true CREs and the predicted CREs.

### Baseline methods

In this study, we compared CREATE to multiple baseline methods by expanding them into multi-class models, including DeepSEA^17^, DanQ^18^, ES-transition^19^ and DeepICSH^20^. DeepSEA was implemented from their original source code (https://deepsea.princeton.edu/). DanQ was implemented from their original source code repository (https://github.com/uci-cbcl/DanQ). ES-transition was implemented from their original source code repository (https://github.com/ncbi/SilencerEnhancerPredict). DeepICSH was implemented from their original source code repository (https://github.com/lyli1013/DeepICSH).

## Statistics and reproducibility

No statistical method was used to predetermine sample size. No data were excluded from the analyses. The experiments were not randomized. Data collection and analysis were not performed blind to the conditions of the experiments.

## Data availability

All datasets used in this study were obtained from public sources. We downloaded experimentally validated silencers from the SilencerDB database^88^ (http://health.tsinghua.edu.cn/SilencerDB/), enhancers from the FANTOM5 project^24, 26^ (https://bioinfo.vanderbilt.edu/AE/HACER/), TSSs from the EPD database^89^ (https://epd.expasy.org/epd), insulators from the ENCODE project^92, 93^ (https://www.encodeproject.org/files/ENCFF085HTY/; https://www.encodeproject.org/files/ENCFF237OKO/) for the K562 and HepG2 cell types. We downloaded the histone modification ChIP-seq peaks and chromatin accessibility peaks from the Roadmap project^94^ (http://egg2.wustl.edu/roadmap/data/byFileType/peaks/consolidated/narrowPeak) and ENCODE project (https://www.encodeproject.org/files/ENCFF055NNT/; https://www.encodeproject.org/files/ENCFF333TAT/; https://www.encodeproject.org/files/ENCFF558BLC/; https://www.encodeproject.org/files/ENCFF842UZU/; https://www.encodeproject.org/files/ENCFF439EIO/; https://www.encodeproject.org/files/ENCFF913MQB/) for the K562 and HepG2 cell types. The non-coding genes were obtained from GENCODE^114^ for human (GRCh37.p13/hg19). All regions in this study are either in the genome of GRCh37/hg19 or have been converted to GRCh37/hg19 by UCSC liftOver^115^ tool.

### Code availability

The CREATE software, including detailed documents and tutorial, is freely available on GitHub (https://github.com/cuixj19/CREATE).

## Acknowledgements

This work was supported by the National Key Research and Development Program of China grant nos. 2021YFF1200902 (R.J.), 2023YFF1204802 (R.J.), the National Natural Science Foundation of China grants no. 62273194 (R.J.).

## Author contributions

R.J. and W.Z. conceived the study and supervised the project. X.C. designed, implemented and validated CREATE. Q.Y., Z.G., Z.L., X.C., S.C. and Q.L. helped analyze the results. X.C. and W.Z. wrote the manuscript, with input from all the authors.

