## Supplementary information for "CREATE: cell-type-specific cis-regulatory elements identification via discrete embedding"

#### Supplementary Figures

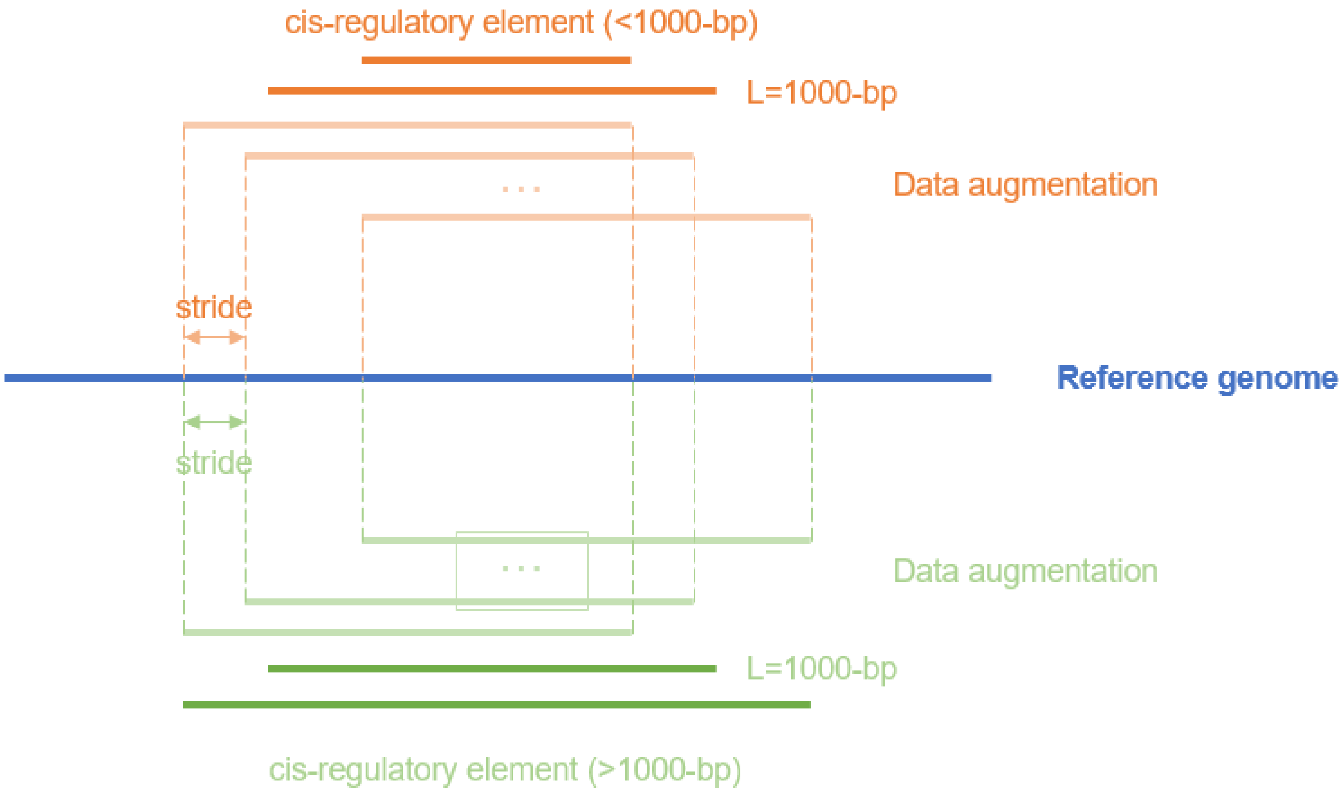

**Supplementary Fig. 1. Data augmentation**. To address the issue of variable length, we adjusted the length of each CRE by extending or reducing it to encompass 1000 base pairs around its midpoint. We slid a window along the genome with stride s (s=10) around the original sequence to obtain augmented sequences.

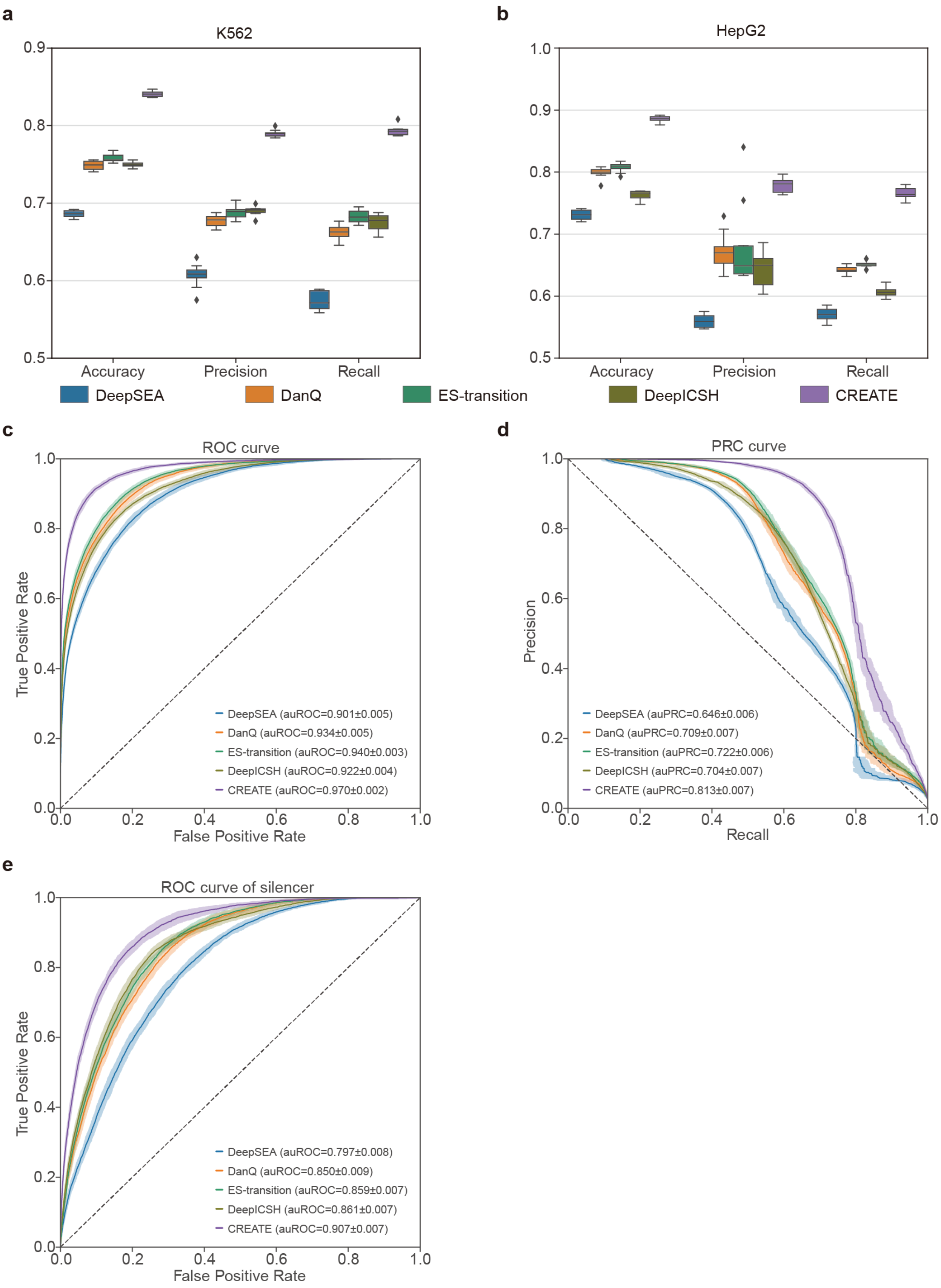

**Supplementary Fig. 2. Evaluation of CREATE compared with the baseline methods**. **a-b**, Boxplot of classification performance evaluated by accuracy, precision and recall on the K562 cell type (**a**) and HepG2 cell type (**b**). Each box plot ranges from the upper to lower quartiles with the median as the horizontal line, whiskers extend to 1.5 times the interquartile range, and points represent outliers. **c-d**, Receiver Operating Characteristic curve (**c**) and Precision-Recall curve (**d**) comparing CREATE and baseline methods on the HepG2 cell type. **e**, ROC curve comparing CREATE and baseline methods for silencers in the K562 cell type. The mean and standard error of auROC or auPRC are reported in the legend. The confidence band shows ±1 s.d. for the averaged curve.

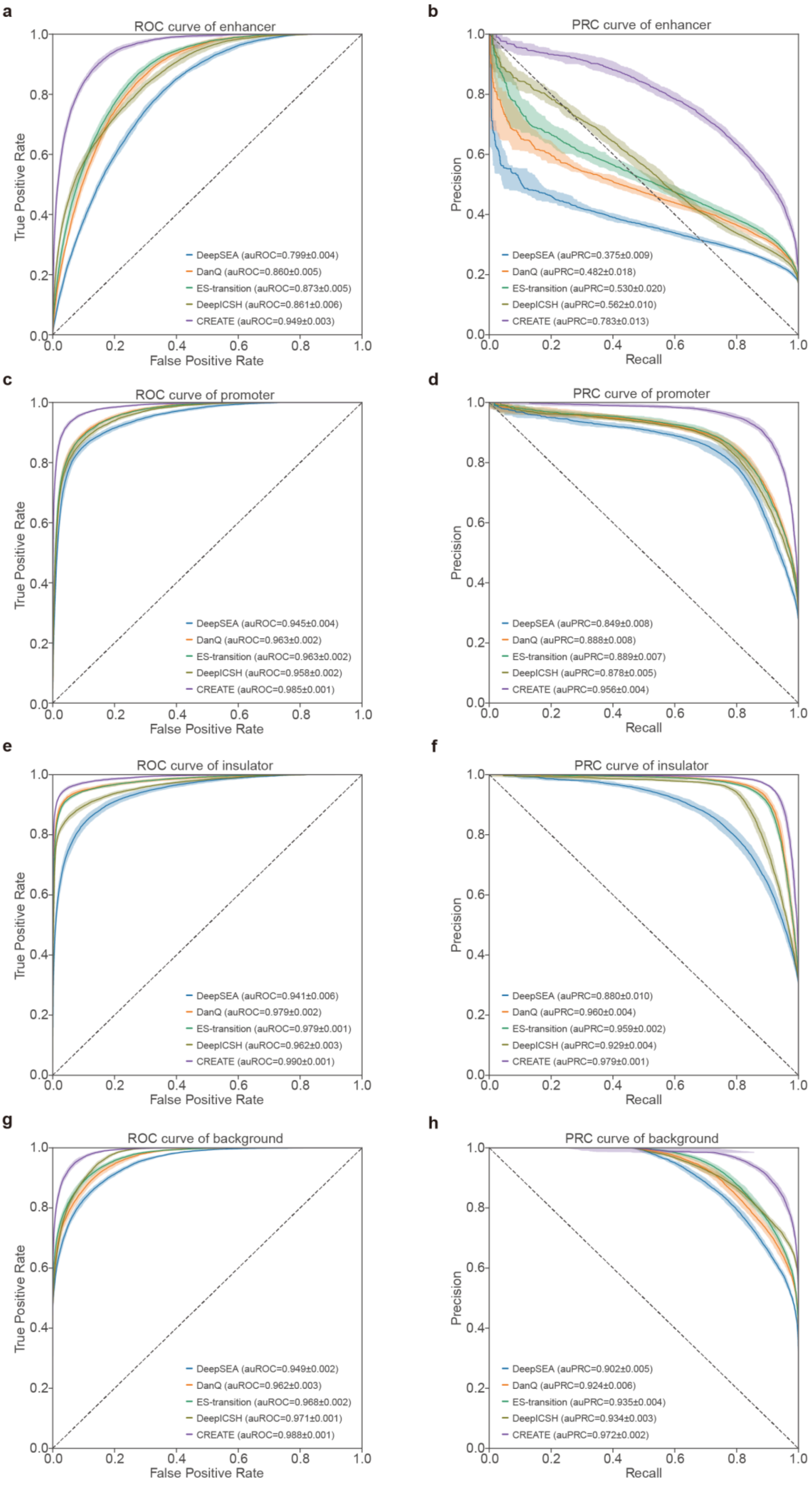

**Supplementary Fig. 3. Evaluation of CREATE compared with the baseline methods**. **a-h**, Receiver Operating Characteristic curve (**a,c,e,g**) and Precision-Recall curve (**b,d,f,h**) curve comparing CREATE and baseline methods for enhancer (**a,b**), promoter (**c,d**), insulator (**e,f**) and background regions (**g,h**) in the K562 cell type. The mean and standard error of auROC or auPRC are reported in the legend. The confidence band shows ±1 s.d. for the averaged curve.

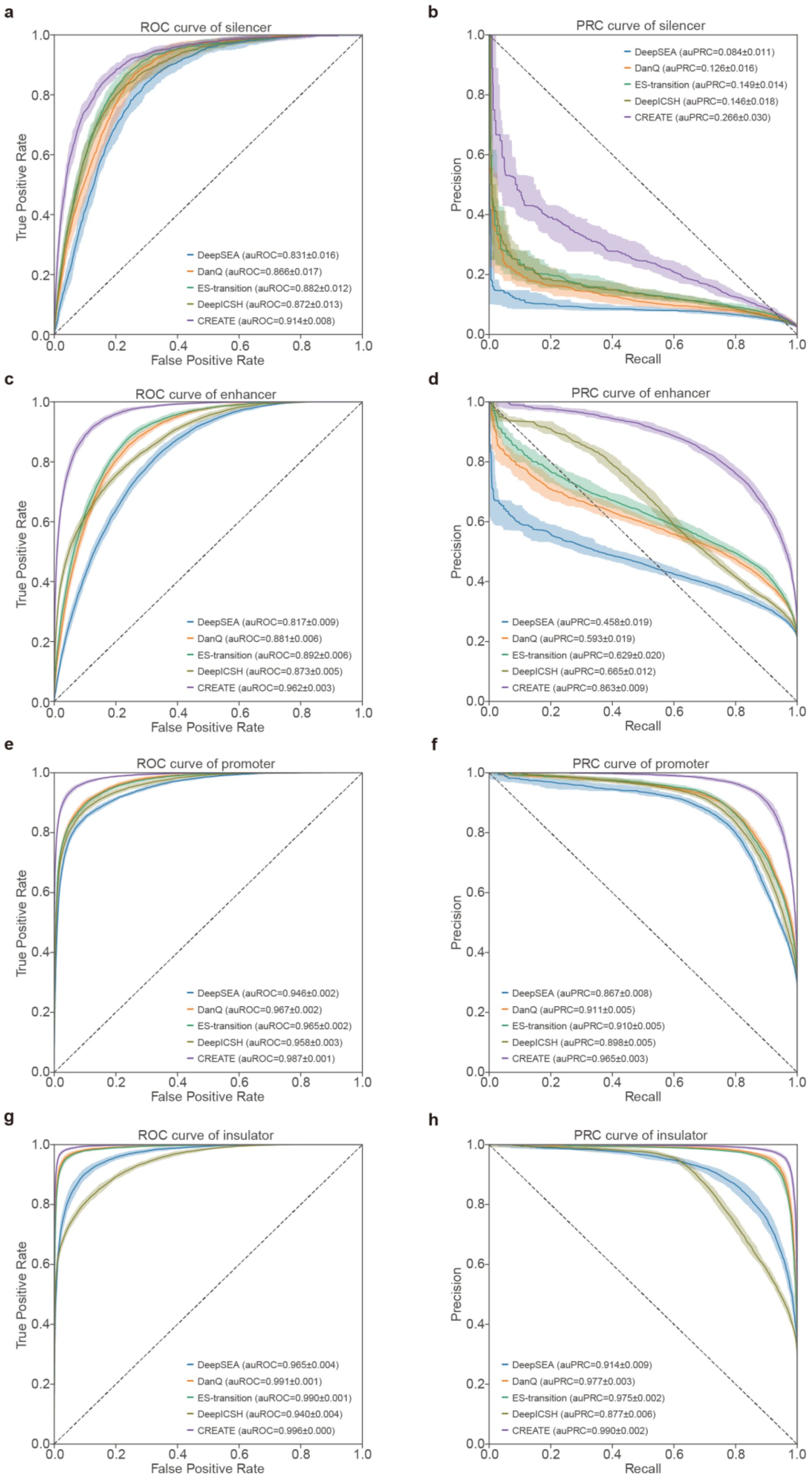

**Supplementary Fig. 4. Evaluation of CREATE compared with the baseline methods**. **a-h**, Receiver Operating Characteristic curve (**a,c,e,g,i**) and Precision-Recall curve (**b,d,f,h,j**) comparing CREATE and baseline methods for silencer (**a,b**), enhancer (**c,d**), promoter (**e,f**), insulator (**g,h**) and background regions (**i,j**) of HepG2 cell type. The mean and standard error of auROC or auPRC are reported in the legend. The confidence band shows ±1 s.d. for the averaged curve.

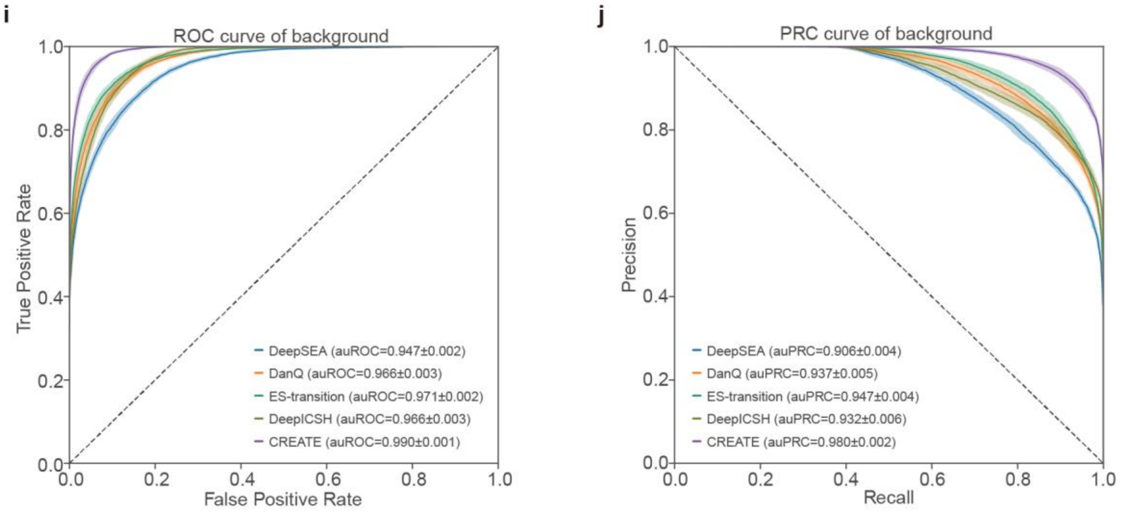

**Supplementary Fig. 4 (continue). Evaluation of CREATE compared with the baseline methods**. **a-h**, Receiver Operating Characteristic curve (**a,c,e,g,i**) and Precision-Recall curve (**b,d,f,h,j**) comparing CREATE and baseline methods for silencer (**a,b**), enhancer (**c,d**), promoter (**e,f**), insulator (**g,h**) and background regions (**i,j**) of HepG2 cell type. The mean and standard error of auROC or auPRC are reported in the legend. The confidence band shows ±1 s.d. for the averaged curve.

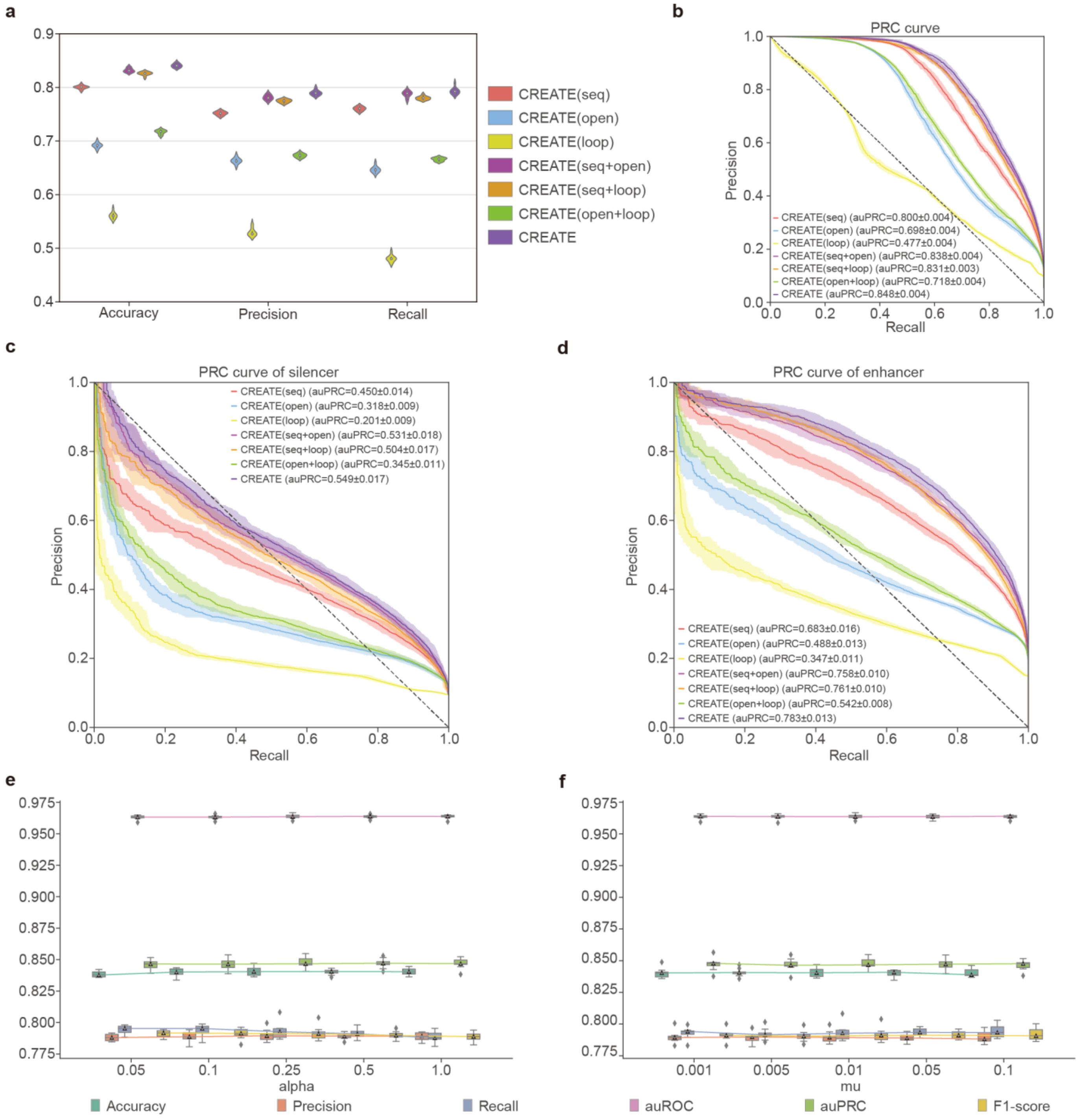

**Supplementary Fig. 5. Robustness analysis of CREATE.** **a**, Violin plot of classification performance evaluated by accuracy, precision and recall for model ablation of CREATE on the K562 cell type. **b**, Precision-Recall curve for model ablation of CREATE on the K562 cell type. **c-d**, Precision-Recall curve for silencers (**c**) and enhancers (**d**) in the K562 cell type for model ablation of CREATE. The mean and standard error of auROC or auPRC are reported in the legend. The confidence band shows ±1 s.d. for the averaged curve. **e**, Classification performance of CREATE under different values of alpha (weight of encoder loss) on the K562 cell type. **f**, Classification performance of CREATE under different values of mu (update ratio of codebook) on the K562 cell type. Each box plot ranges from the upper to lower quartiles with the median as the horizontal line, whiskers extend to 1.5 times the interquartile range, and points represent outliers.

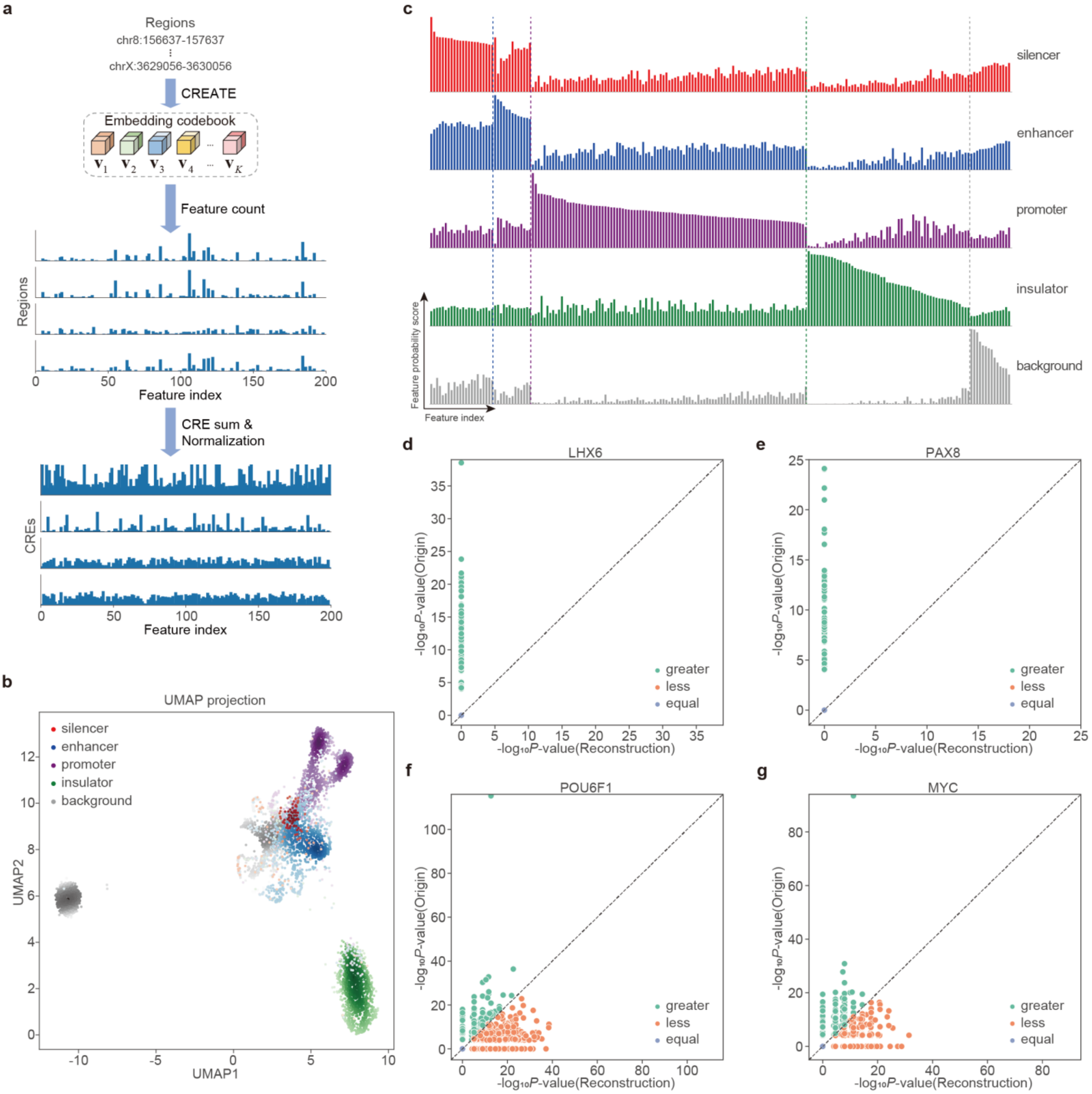

**Supplementary Fig. 6. Generation and interpretation of CRE-specific feature spectrum. a**, Feature count matrix, in which rows correspond to CRE regions and columns to the feature indices in the codebook, is extracted, summed along CREs, normalized and reordered to form a feature spectrum for each CRE. **b**, UMAP visualization of the CRE embeddings from CREATE on the testing data in one of the 10-fold cross-validation experiments of HepG2 cell type. **c**, CRE-specific feature spectrum. There are a distinct set of specific features that are enriched or depleted in the feature spectrum of each CRE on the HepG2 cell type. **d-g**, Comparison of LHX6 (**d**), PAX8 (**e**), POU6F1 (**f**) and MYC (**g**) motif enrichment significance (-$\log_{10}$*P*-value) between original input and reconstructed output when information derived from the major feature in the silencer-specific feature spectrum of K562 cell type is removed by zeroing it out before passing the CRE embeddings again through the decoder.

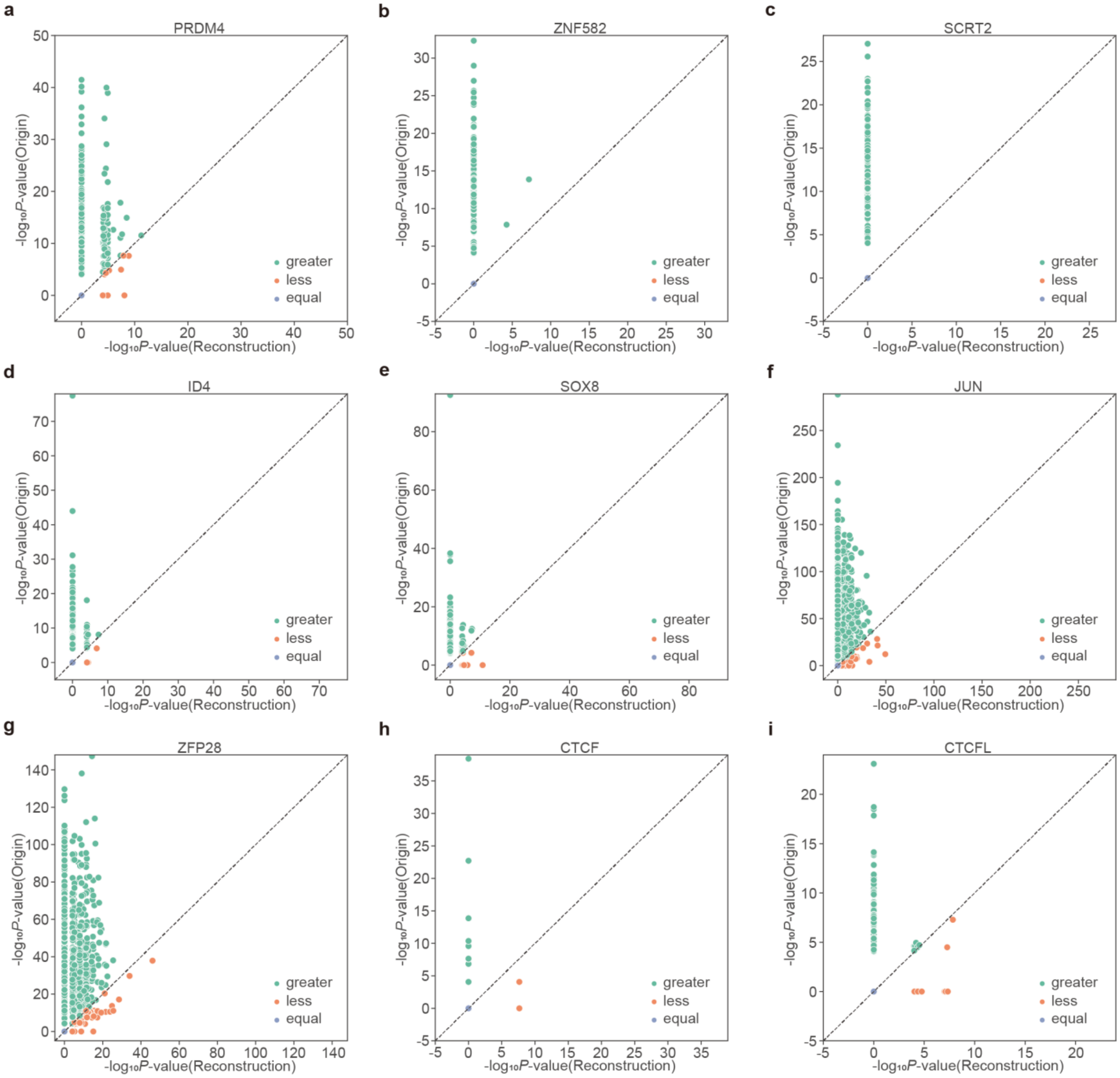

**Supplementary Fig. 7. Comparison of motif enrichment significance between original input and reconstructed output. a-c**, Comparison of PRDM4 (**a**), ZNF582 (**b**) and SCRT2 (**c**) motif enrichment significance (-log$_{10}$*P*-value) between original input and reconstructed output when information derived from the major feature in the silencer-specific feature spectrum of K562 cell type is removed by zeroing it out before passing the CRE embeddings again through the decoder. **d-e**, Comparison of ID4 (**d**) and SOX8 (**e**) motif enrichment significance (-log$_{10}$*P*-value) between original input and reconstructed output when information derived from the major feature in the enhancer-specific feature spectrum of K562 cell type is removed. **f-g**, Comparison of JUN (**f**) and ZFP28 (**g**) motif enrichment significance (-log$_{10}$*P*-value) between original input and reconstructed output when information derived from the major feature in the promoter-specific feature spectrum of K562 cell type is removed. **h-i**, Comparison of CTCF (**h**) and CTCFL (**i**) motif enrichment significance (-log$_{10}$*P*-value) between original input and reconstructed output when information derived from the major feature in the insulator-specific feature spectrum of K562 cell type is removed.

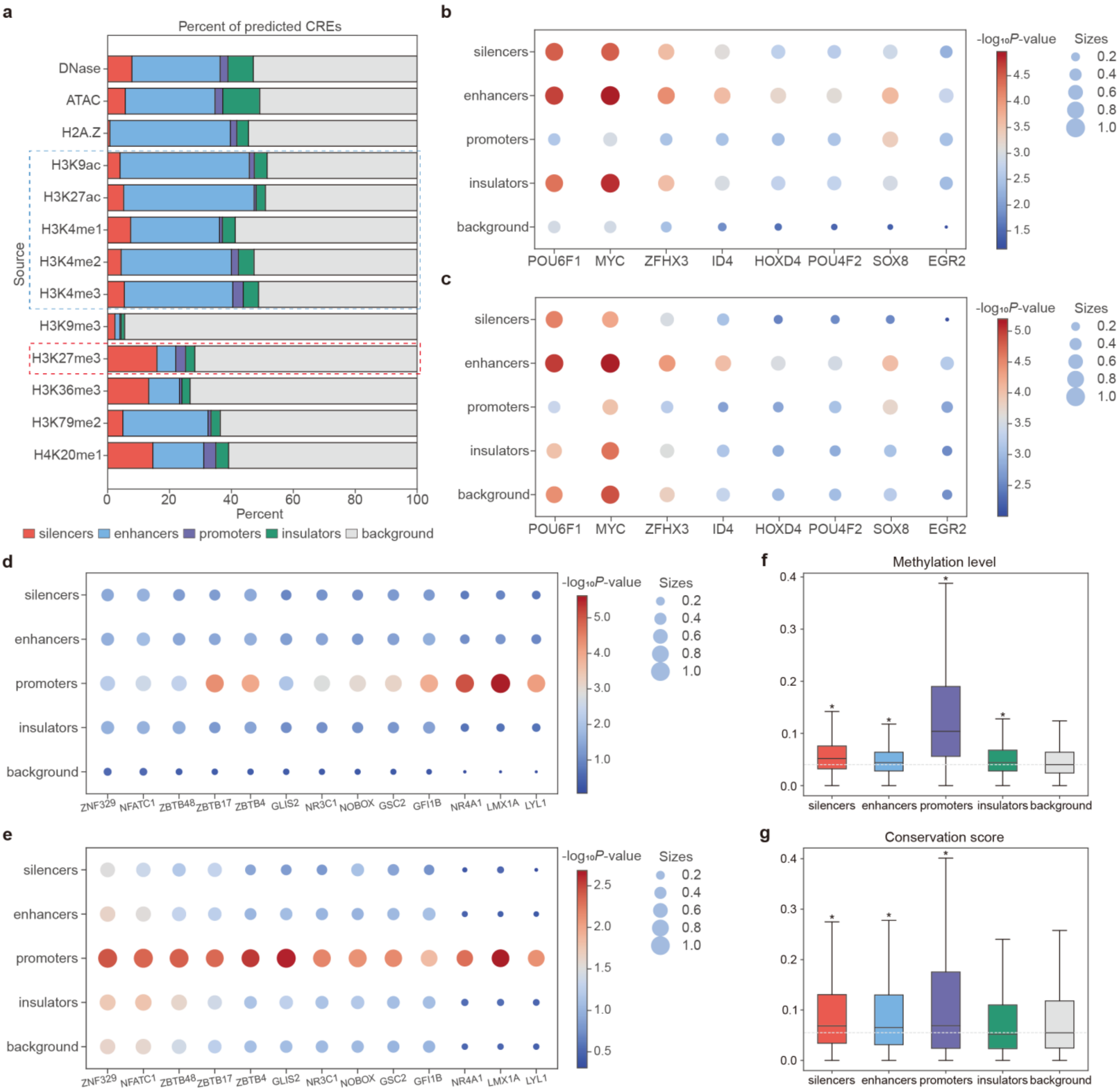

**Supplementary Fig. 8. Characteristics of predicted CREs by CREATE. a**, Percentage of predicted CREs from different candidate sources in the HepG2 cell type. **b-c**, Bubble plot of motif enrichment significance (-log$_{10}$*P*-value) of active TFs (enhancer-related TFs) at true CREs (**b**) and predicted CREs (**c**) on the K562 cell type. **d-e**, Bubble plot of motif enrichment significance (-log$_{10}$*P*-value) of promoter-related TFs at true CREs (**d**) and predicted CREs (**e**) on the K562 cell type. The size of bubbles represents the proportion of CREs with *P*-value < 0.01. **f**, Box plot of methylation levels at predicted CREs and background regions on the K562 cell type. The asterisks above the boxes indicate the significant enrichments compared with the background regions. (*) *P*-value < 2e-6. **g**, Box plot of conservation scores at predicted CREs and background regions on the K562 cell type. (*) *P*-value < 1e-13. Each box plot ranges from the upper to lower quartiles with the median as the horizontal line, whiskers extend to 1.5 times the interquartile range.

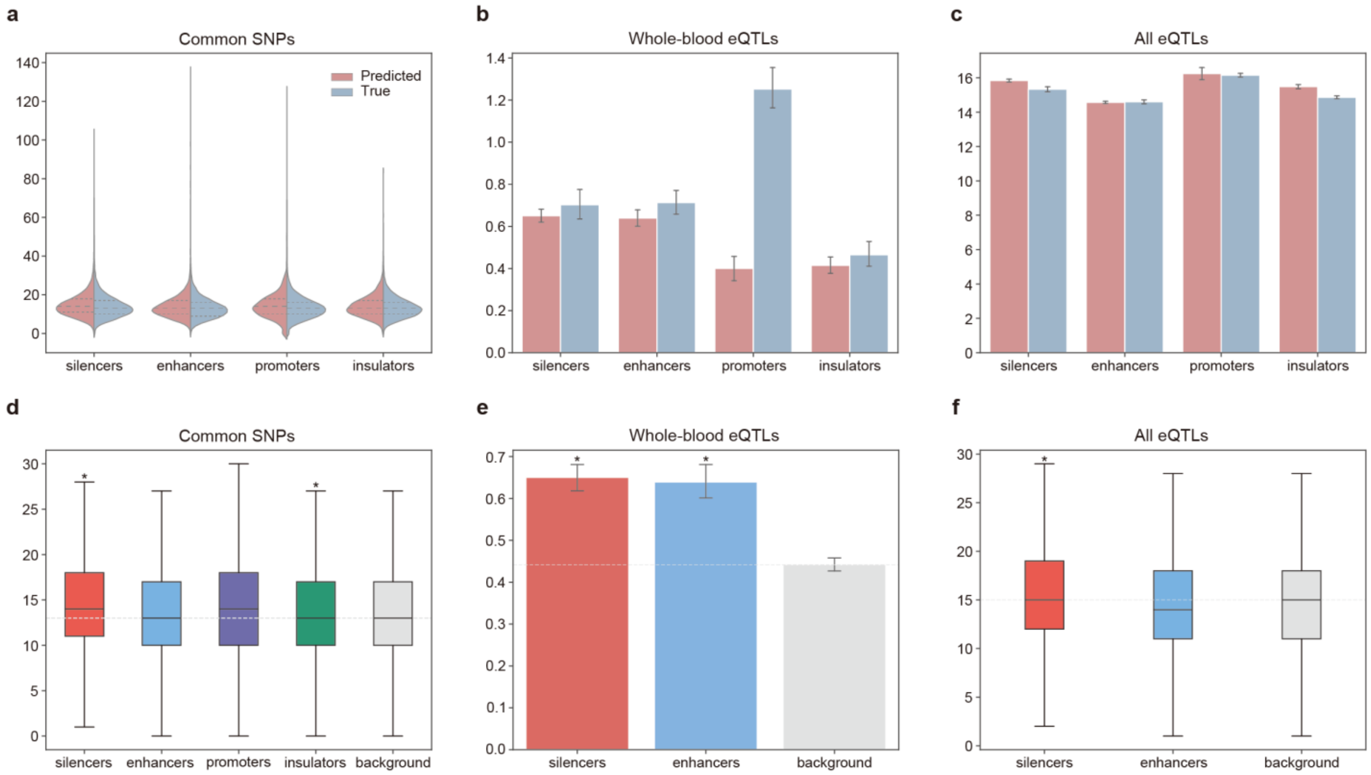

**Supplementary Fig. 9. Identification of the biological variability of CREs by CREATE. a**, Violin plot of overlaps between the common SNPs and true CREs or predicted CREs on the K562 cell type. Each violin plot contains three horizontal dashed lines denoting the median, the upper quartile, and the lower quartile. **b**, Bar plot of overlaps between the whole-blood eQTLs and true CREs or predicted CREs on the K562 cell type. **c**, Bar plot of overlaps between all GTEx eQTLs and true CREs or predicted CREs on the K562 cell type. The error bars denote the 95% confidence interval, and the centers of error bars denote the average value. **d**, Box plot of overlaps between the common SNPs and the predicted CREs or background regions on the K562 cell type. The asterisks above the boxes indicate the significant enrichments compared with the background regions. (∗) $P$-value < 5e-3. Each box plot ranges from the upper to lower quartiles with the median as the horizontal line, whiskers extend to 1.5 times the interquartile range. **e**, Bar plot of overlaps between the whole-blood eQTLs and the predicted silencers, enhancers or background regions on the K562 cell type. (∗) $P$-value < 4e-49. **f**, Bar plot of overlaps between all GTEx eQTLs and the predicted silencers, enhancers or background regions on the K562 cell type. (∗) $P$-value < 2e-28.

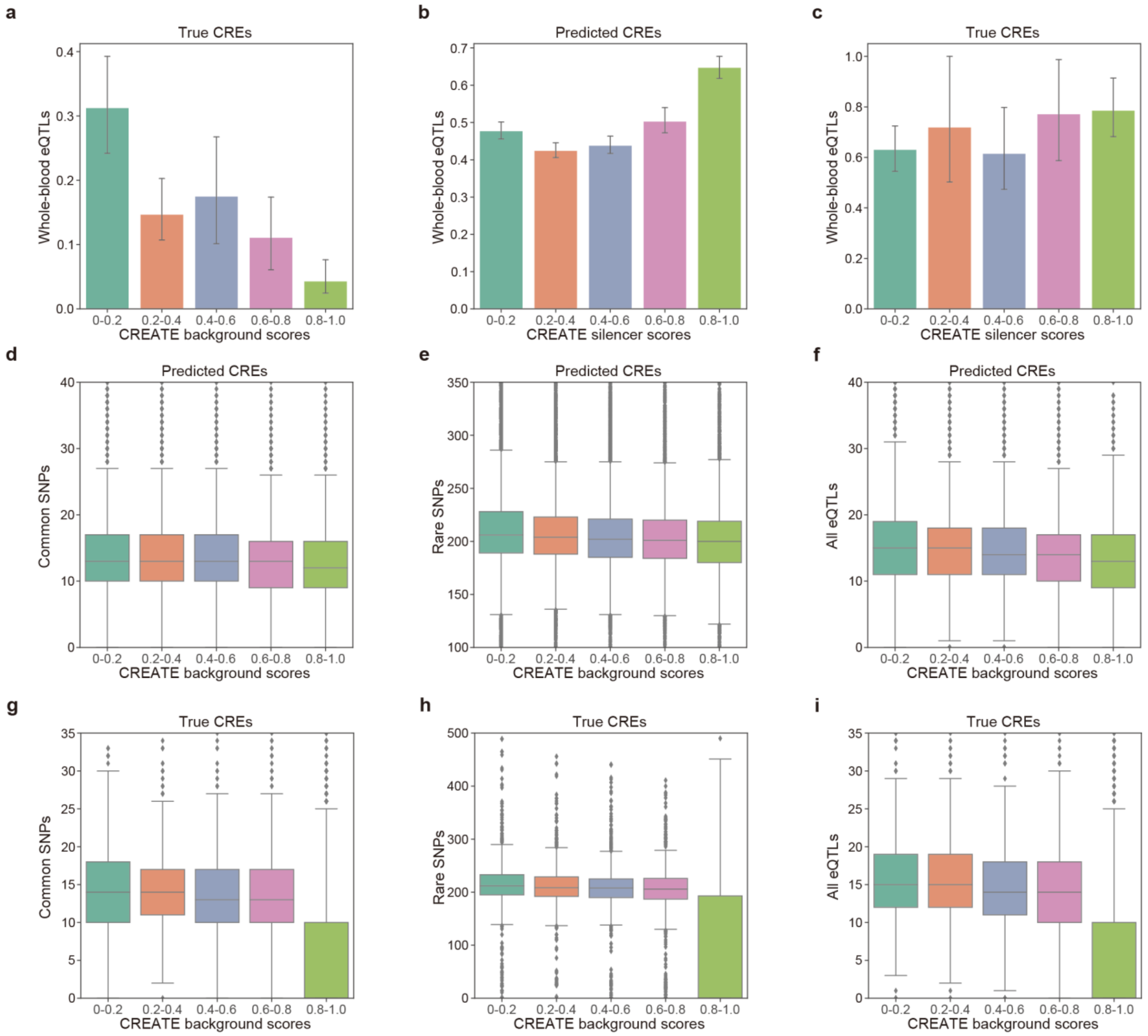

**Supplementary Fig. 10. Identification of the biological variability of CREs by CREATE. a-i**, Correlation between the CREATE background scores (**a**,**d-i**) or CREATE silencer scores (**b-c**) and overlaps between the whole-blood eQTLs (**a-c**), common SNPs (**d**,**g**), rare SNPs (**e**,**h**) or all GTEx eQTLs (**f**,**i**) and true CREs (**a**,**c**,**g**,**h**,**i**) or predicted CREs (**b**,**d**,**e**,**f**) on the K562 cell type. The error bars of bar plots denote the 95% confidence interval, and the centers of error bars denote the average value. Each box plot ranges from the upper to lower quartiles with the median as the horizontal line, whiskers extend to 1.5 times the interquartile range, and points represent outliers.

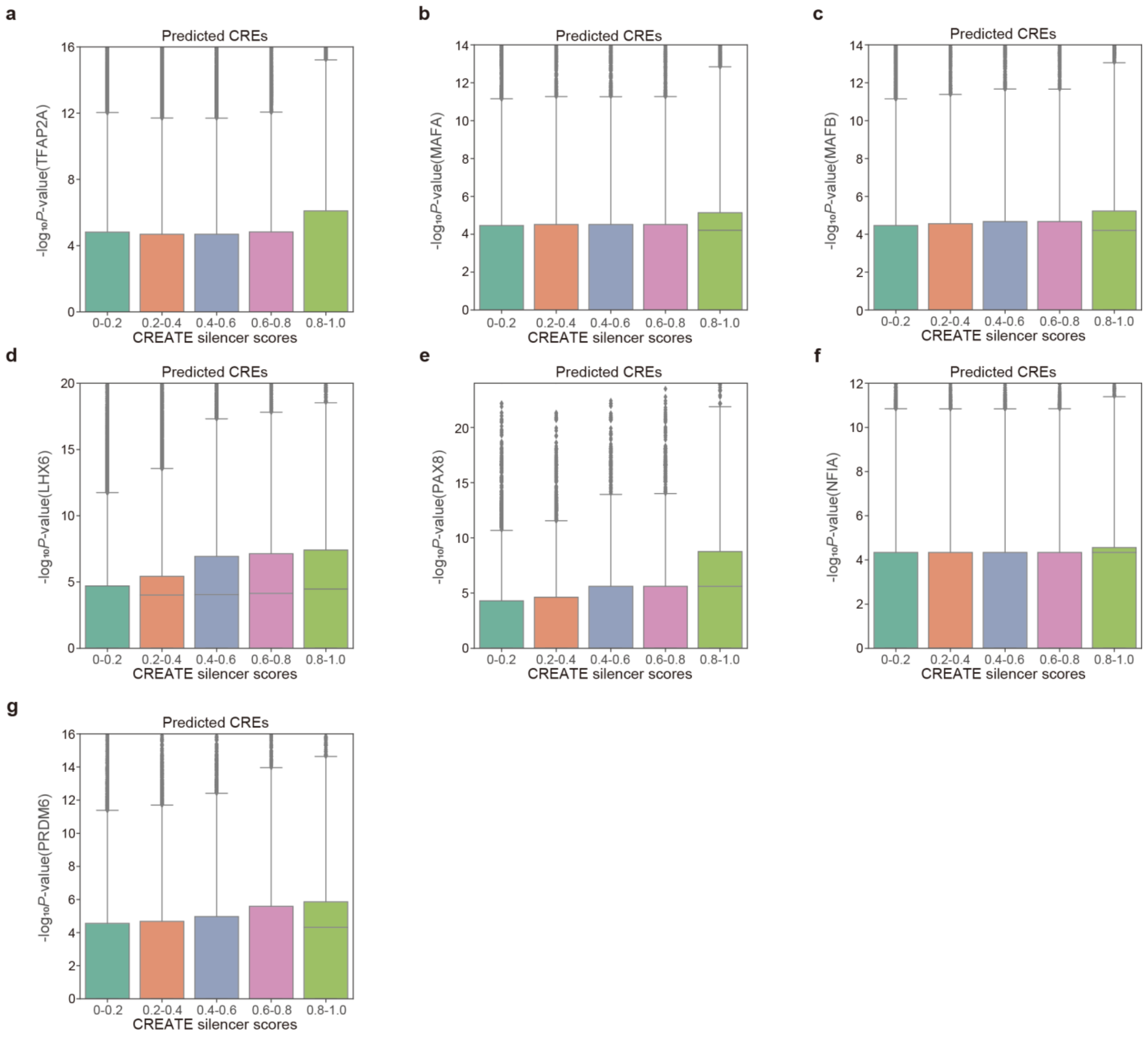

**Supplementary Fig. 11. Correlation between the CREATE silencer scores and the motif enrichment significance. a-g**, Correlation between the CREATE silencer scores and the motif enrichment significance (-$\log_{10}$*P*-value) of TFAP2A (**a**), MAFA (**b**), MAFB (**c**), LHX6 (**d**), PAX8 (**e**), NFIA (**f**) and PRDM6 (**g**) at predicted CREs on the K562 cell type. Each box plot ranges from the upper to lower quartiles with the median as the horizontal line, whiskers extend to 1.5 times the interquartile range, and points represent outliers.

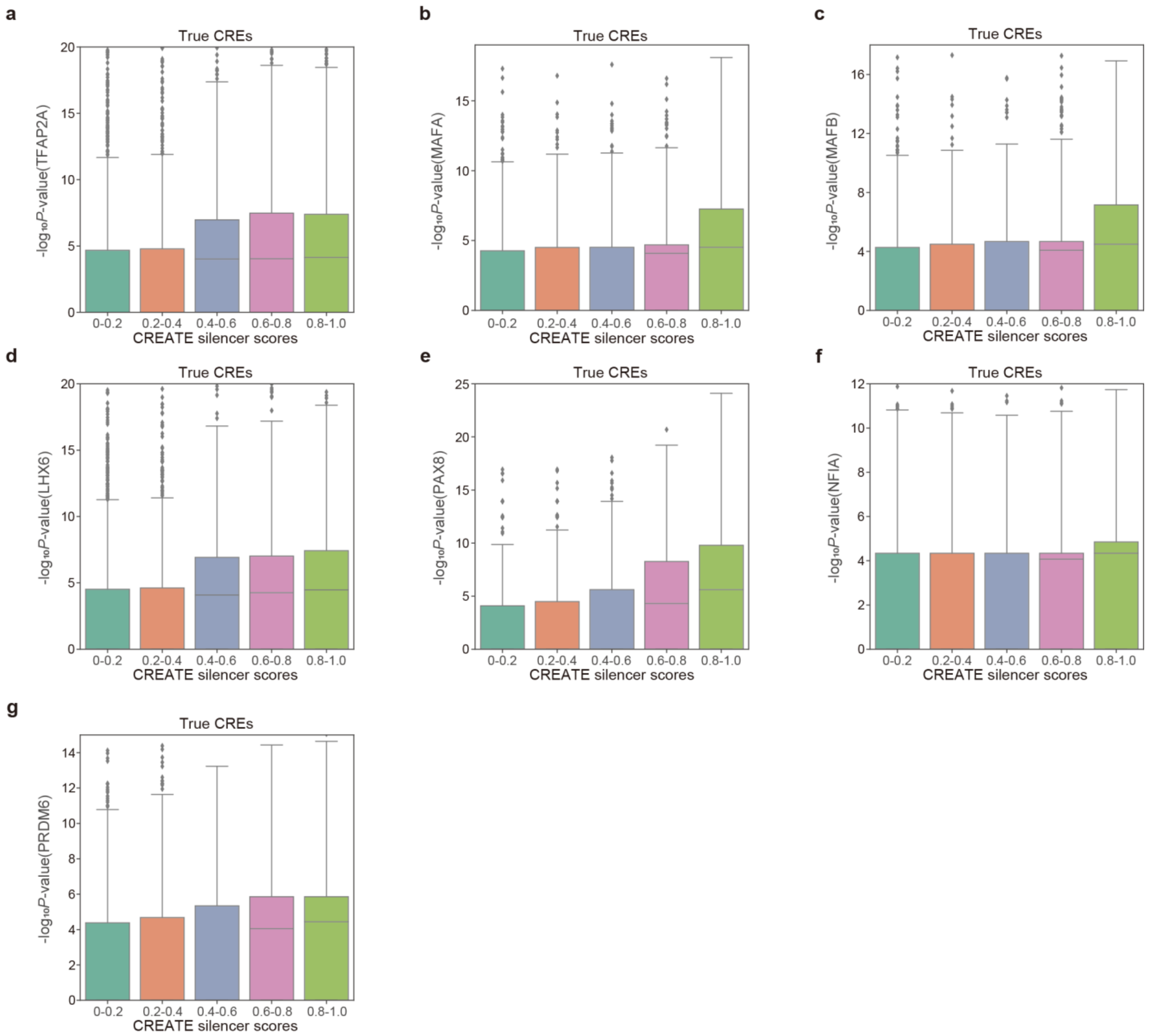

**Supplementary Fig. 12. Correlation between the CREATE silencer scores and the motif enrichment significance. a-g**, Correlation between the CREATE silencer scores and the motif enrichment significance (-log$_{10}$*P*-value) of TFAP2A (**a**), MAFA (**b**), MAFB (**c**), LHX6 (**d**), PAX8 (**e**), NFIA (**f**) and PRDM6 (**g**) at true CREs on the K562 cell type. Each box plot ranges from the upper to lower quartiles with the median as the horizontal line, whiskers extend to 1.5 times the interquartile range, and points represent outliers.

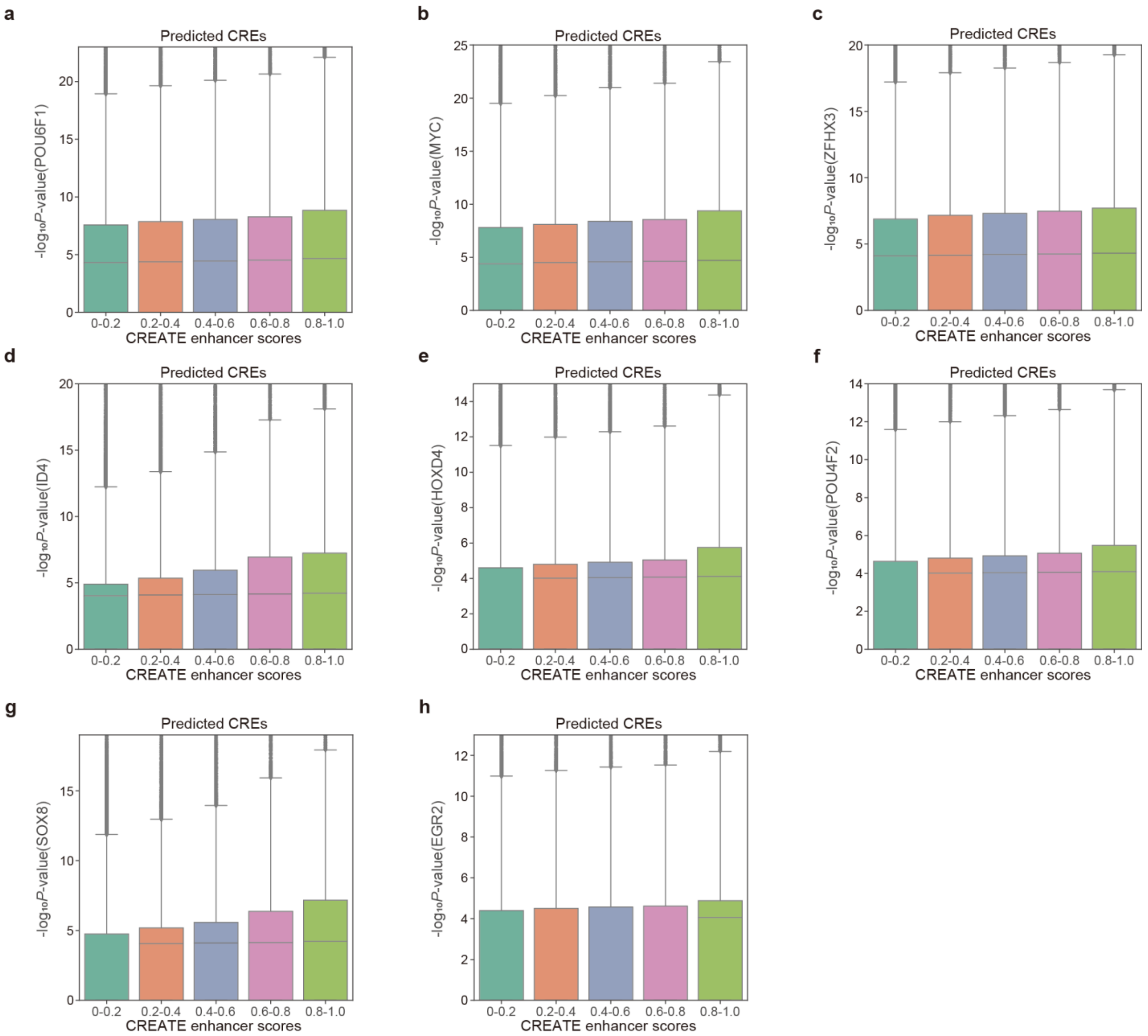

**Supplementary Fig. 13. Correlation between the CREATE enhancer scores and the motif enrichment significance. a-h**, Correlation between the CREATE enhancer scores and the motif enrichment significance (-log$_{10}$*P*-value) of POU6F1 (**a**), MYC (**b**), ZFHX3 (**c**), ID4 (**d**), HOXD4 (**e**), POU4F2 (**f**), SOX8 (**g**) and EGR2 (**h**) at predicted CREs on the K562 cell type. Each box plot ranges from the upper to lower quartiles with the median as the horizontal line, whiskers extend to 1.5 times the interquartile range, and points represent outliers.

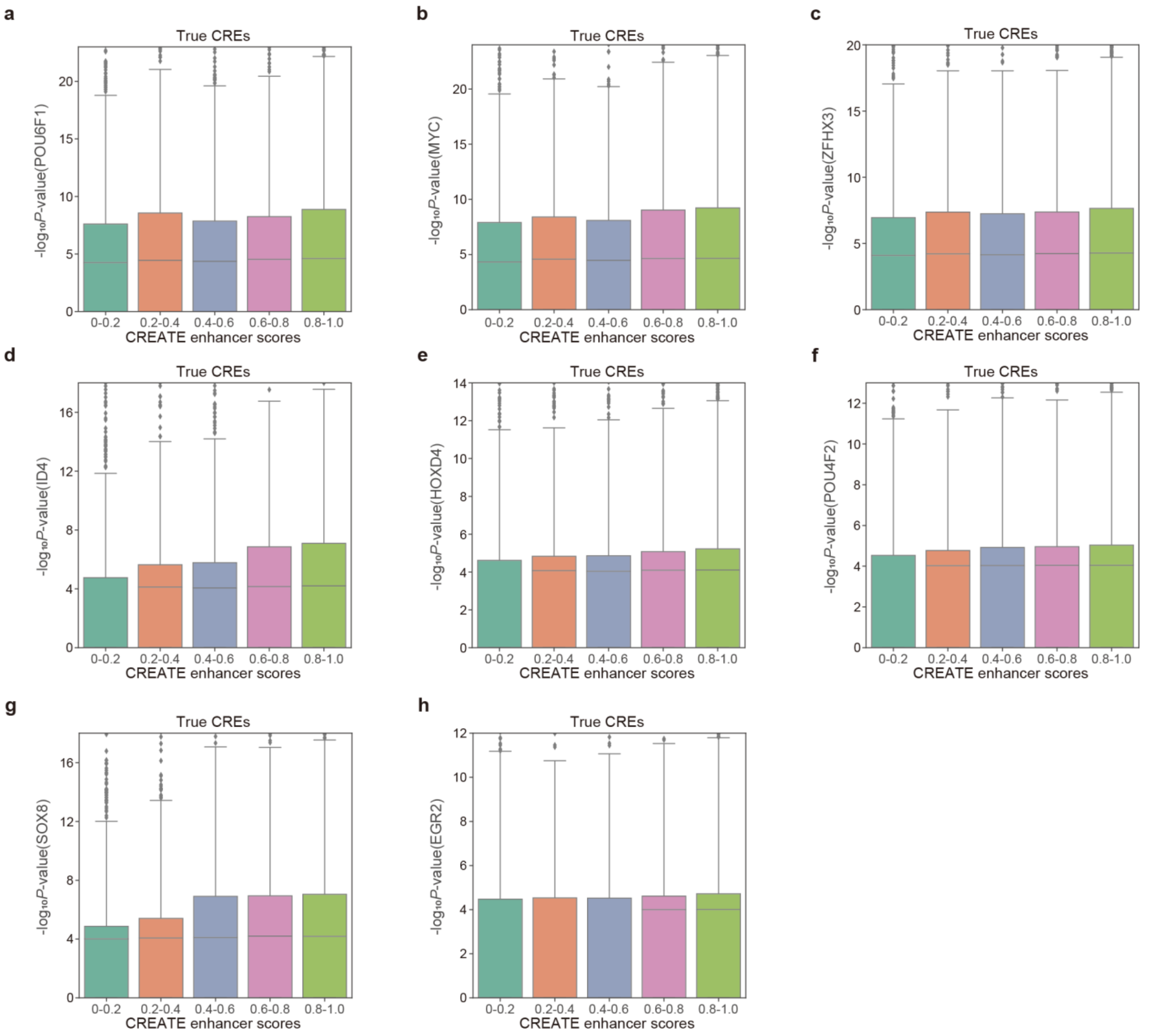

**Supplementary Fig. 14. Correlation between the CREATE enhancer scores and the motif enrichment significance. a-h**, Correlation between the CREATE enhancer scores and the motif enrichment significance (-$\log_{10}P$-value) of POU6F1 (**a**), MYC (**b**), ZFHX3 (**c**), ID4 (**d**), HOXD4 (**e**), POU4F2 (**f**), SOX8 (**g**) and EGR2 (**h**) at true CREs on the K562 cell type. Each box plot ranges from the upper to lower quartiles with the median as the horizontal line, whiskers extend to 1.5 times the interquartile range, and points represent outliers.

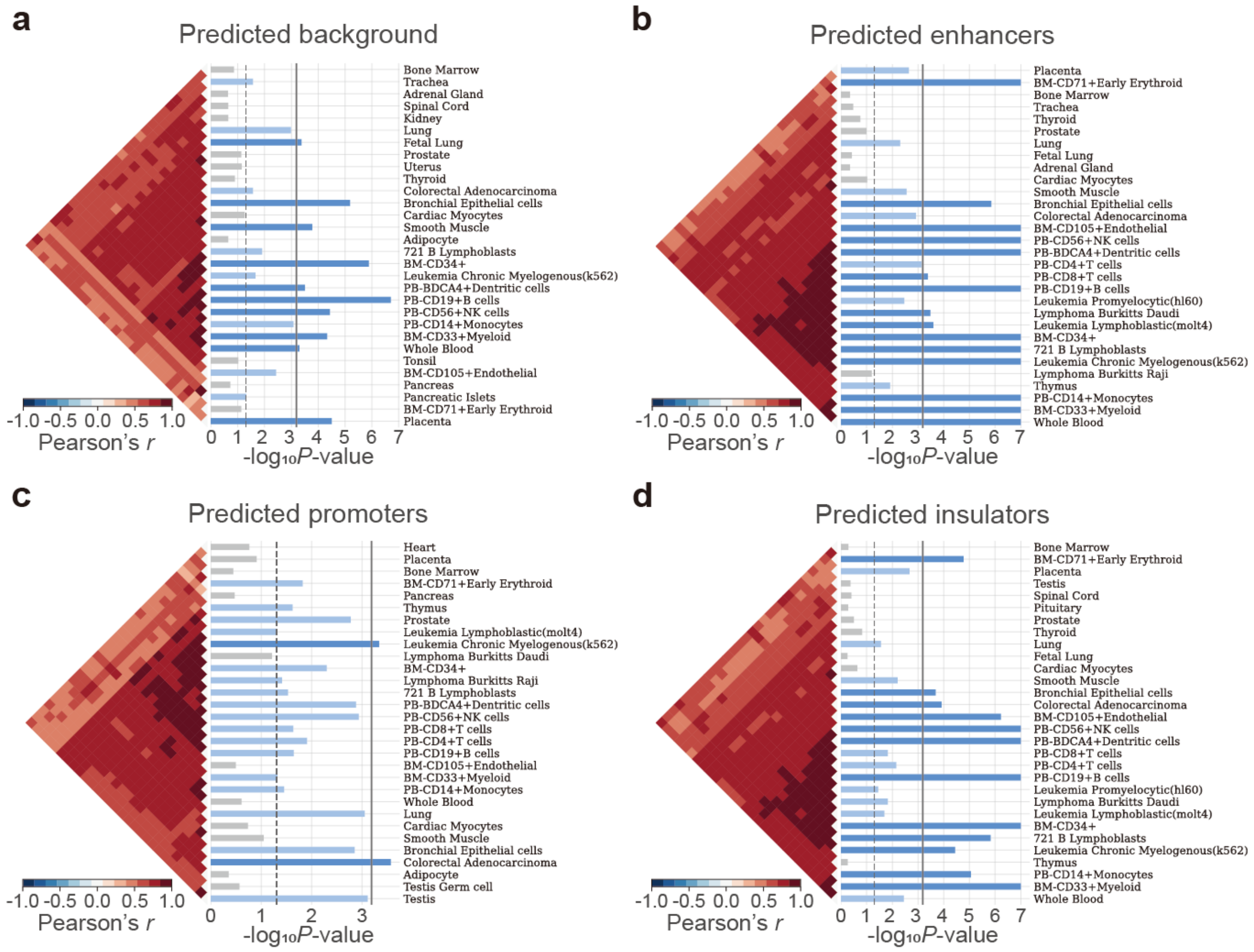

**Supplementary Fig. 15. Tissue enrichment analysis for predicted CREs and background regions on the K562 cell type. a-e**, Top 30 significantly enriched tissues in SNPsea analysis on predicted background regions (**a**), predicted enhancers (**b**), predicted promoters (**c**) and predicted insulators (**d**). The vertical dashed line represents the one-sided *P*-value cutoff at the 0.05 level, while the solid lines denotes the cutoff at 0.05 level for the one-sided *P*-value with Bonferroni correction. Each plot also contains the ordered expression profiles using hierarchical clustering with unweighted pair-group method with arithmetic means, and the Pearson correlation coefficients indicating the correlation between profiles.

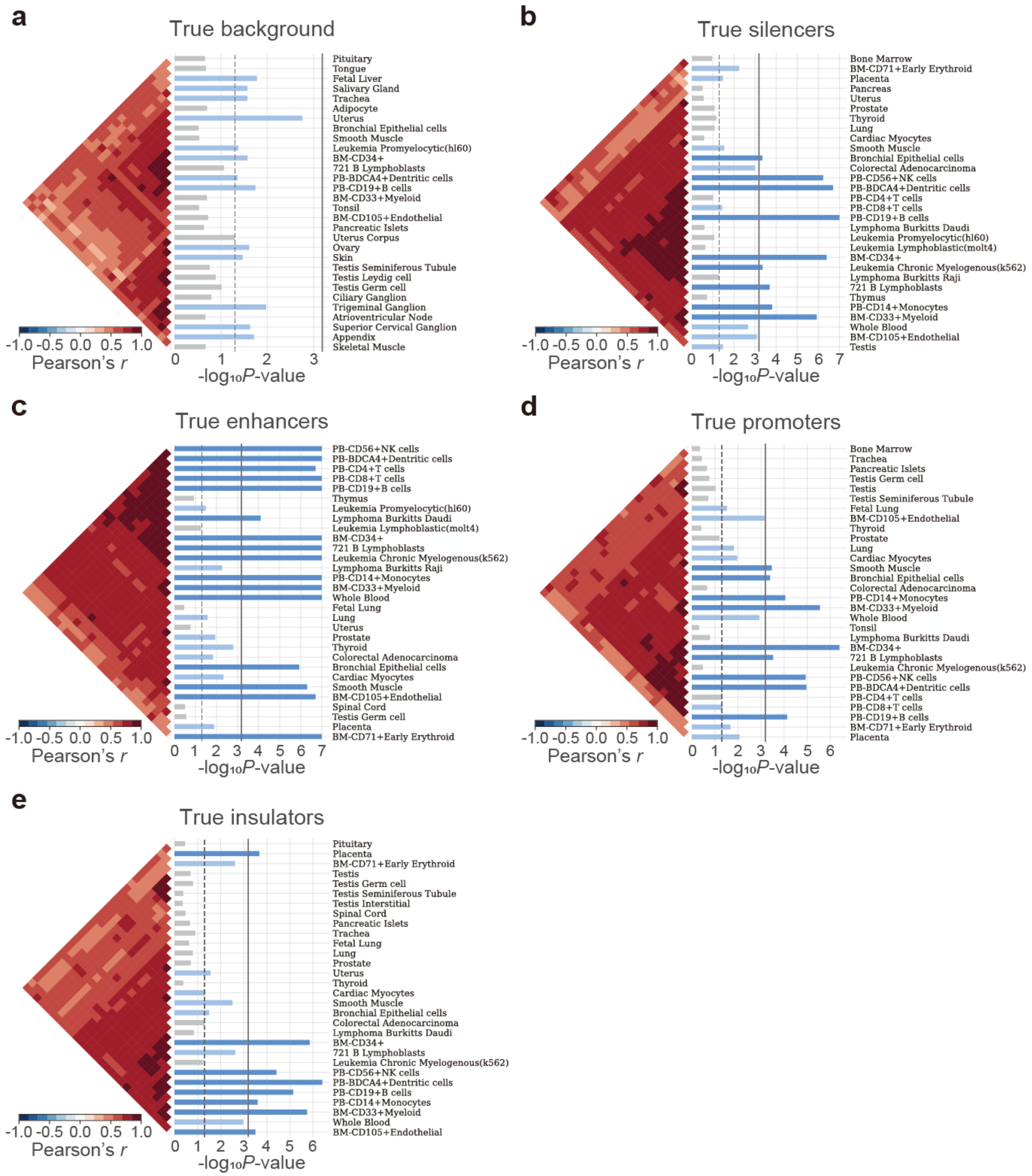

**Supplementary Fig. 16. Tissue enrichment analysis for true CREs and background regions on the K562 cell type. a-e**, Top 30 significantly enriched tissues in SNPsea analysis on true background regions (**a**), true silencers (**b**), true enhancers (**c**), true promoters (**d**) and true insulators (**e**). The vertical dashed line represents the one-sided *P*-value cutoff at the 0.05 level, while the solid lines denotes the cutoff at 0.05 level for the one-sided *P*-value with Bonferroni correction. Each plot also contains the ordered expression profiles using hierarchical clustering with unweighted pair-group method with arithmetic means, and the Pearson correlation coefficients indicating the correlation between profiles.

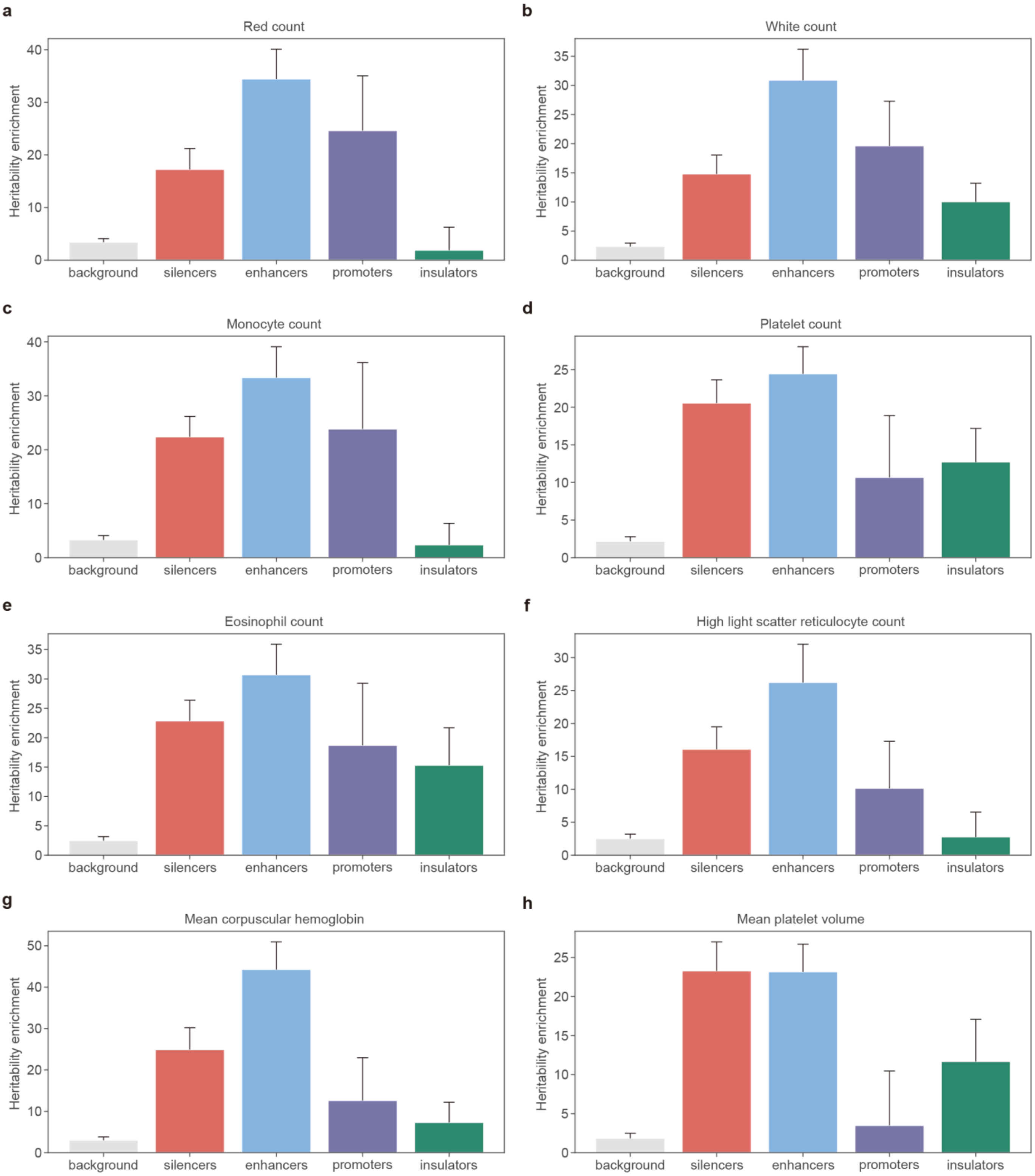

**Supplementary Fig. 17. Heritability enrichment analysis for predicted CREs and background regions on the K562 cell type. a-m**, Heritability enrichments estimated by LDSC within predicted CREs and background regions identified by CREATE for blood-related traits including red count (**a**), white count (**b**), monocyte count (**c**), platelet count (**d**), eosinophil count (**e**), high light scatter reticulocyte count (**f**), mean corpuscular hemoglobin (**g**), mean platelet volume (**h**), mean sphered cell volume (**i**), platelet distribution width (**j**), red blood cell distribution width (**k**), HbA1c (**l**) and albumin (**m**). The error bars denote jackknife standard errors over 200 equally sized blocks of adjacent SNPs about the estimates of enrichment, and the centers of error bars represent the average value.

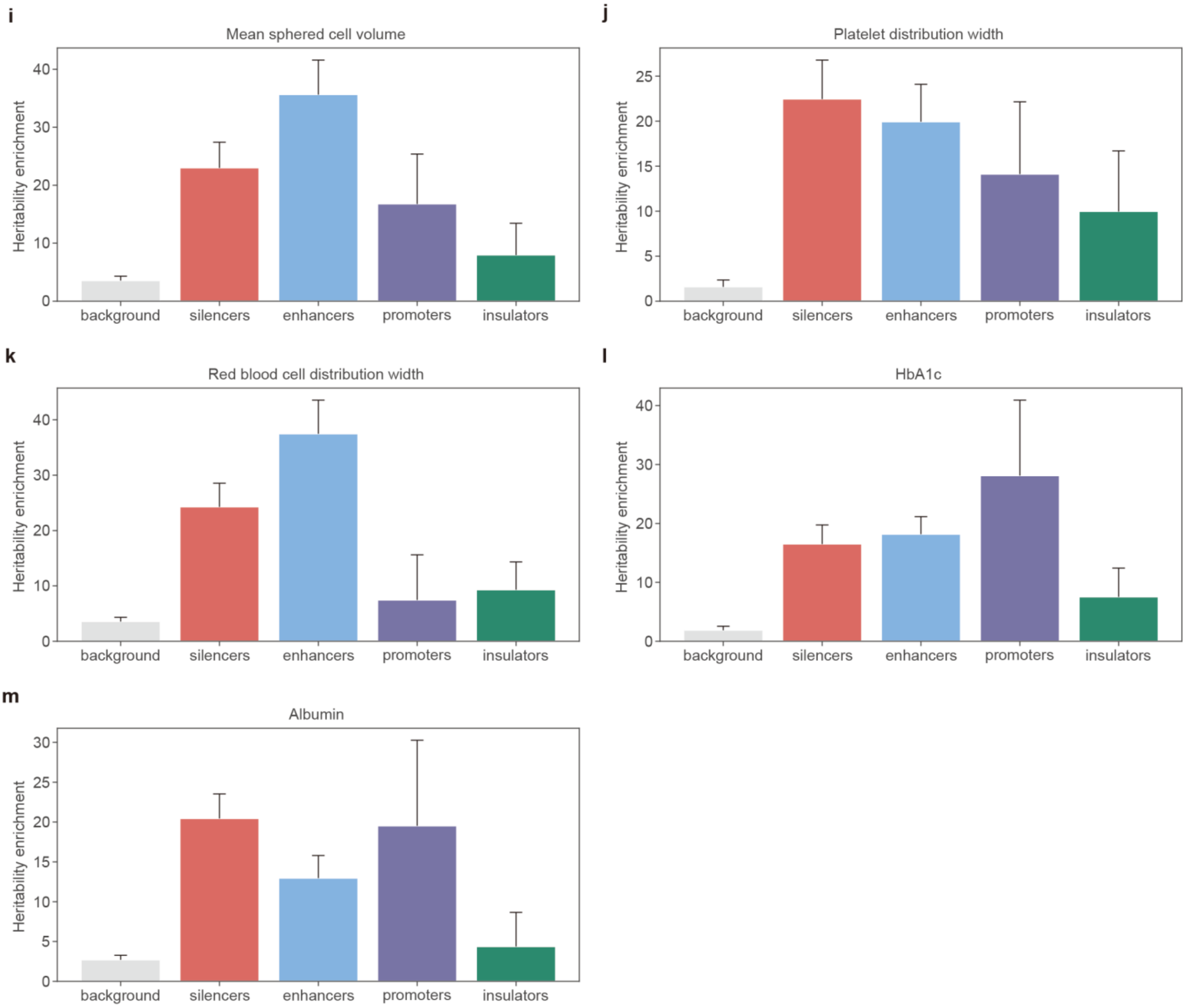

**Supplementary Fig. 17 (continue). Heritability enrichment analysis for predicted CREs and background regions on the K562 cell type. a-m**, Heritability enrichments estimated by LDSC within predicted CREs and background regions identified by CREATE for blood-related traits including red count (**a**), white count (**b**), monocyte count (**c**), platelet count (**d**), eosinophil count (**e**), high light scatter reticulocyte count (**f**), mean corpuscular hemoglobin (**g**), mean platelet volume (**h**), mean sphered cell volume (**i**), platelet distribution width (**j**), red blood cell distribution width (**k**), HbA1c (**l**) and albumin (**m**). The error bars denote jackknife standard errors over 200 equally sized blocks of adjacent SNPs about the estimates of enrichment, and the centers of error bars represent the average value.

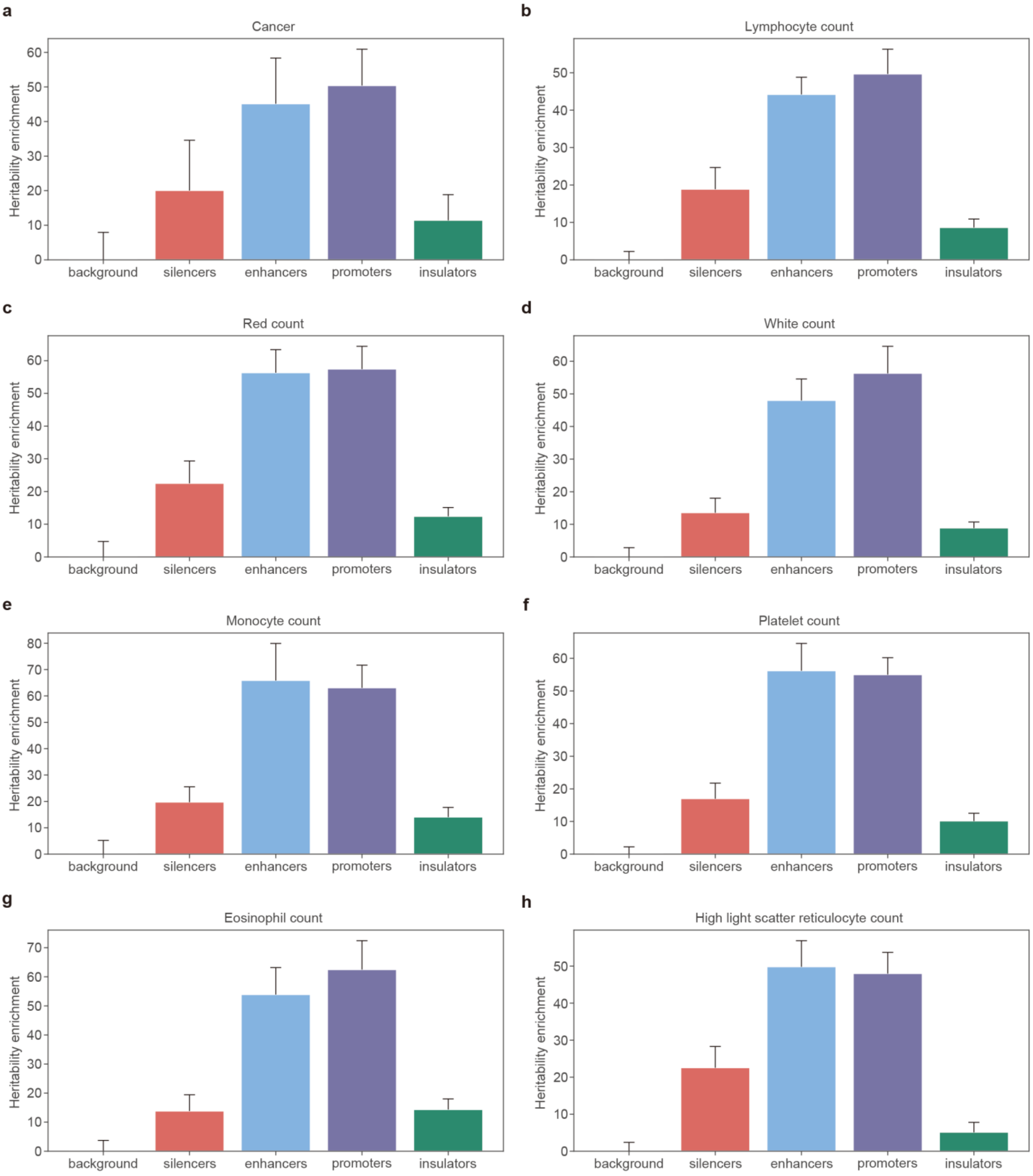

**Supplementary Fig. 18. Heritability enrichment analysis for true CREs and background regions on the K562 cell type. a-o**, Heritability enrichments estimated by LDSC within true CREs and background regions for blood-related traits including cancer (**a**), lymphocyte count (**b**), red count (**c**), white count (**d**), monocyte count (**e**), platelet count (**f**), eosinophil count (**g**), high light scatter reticulocyte count (**h**), mean corpuscular hemoglobin (**i**), mean platelet volume (**j**), mean sphered cell volume (**k**), platelet distribution width (**l**), red blood cell distribution width (**m**), HbA1c (**n**) and albumin (**o**). The error bars denote jackknife standard errors over 200 equally sized blocks of adjacent SNPs about the estimates of enrichment, and the centers of error bars represent the average value.

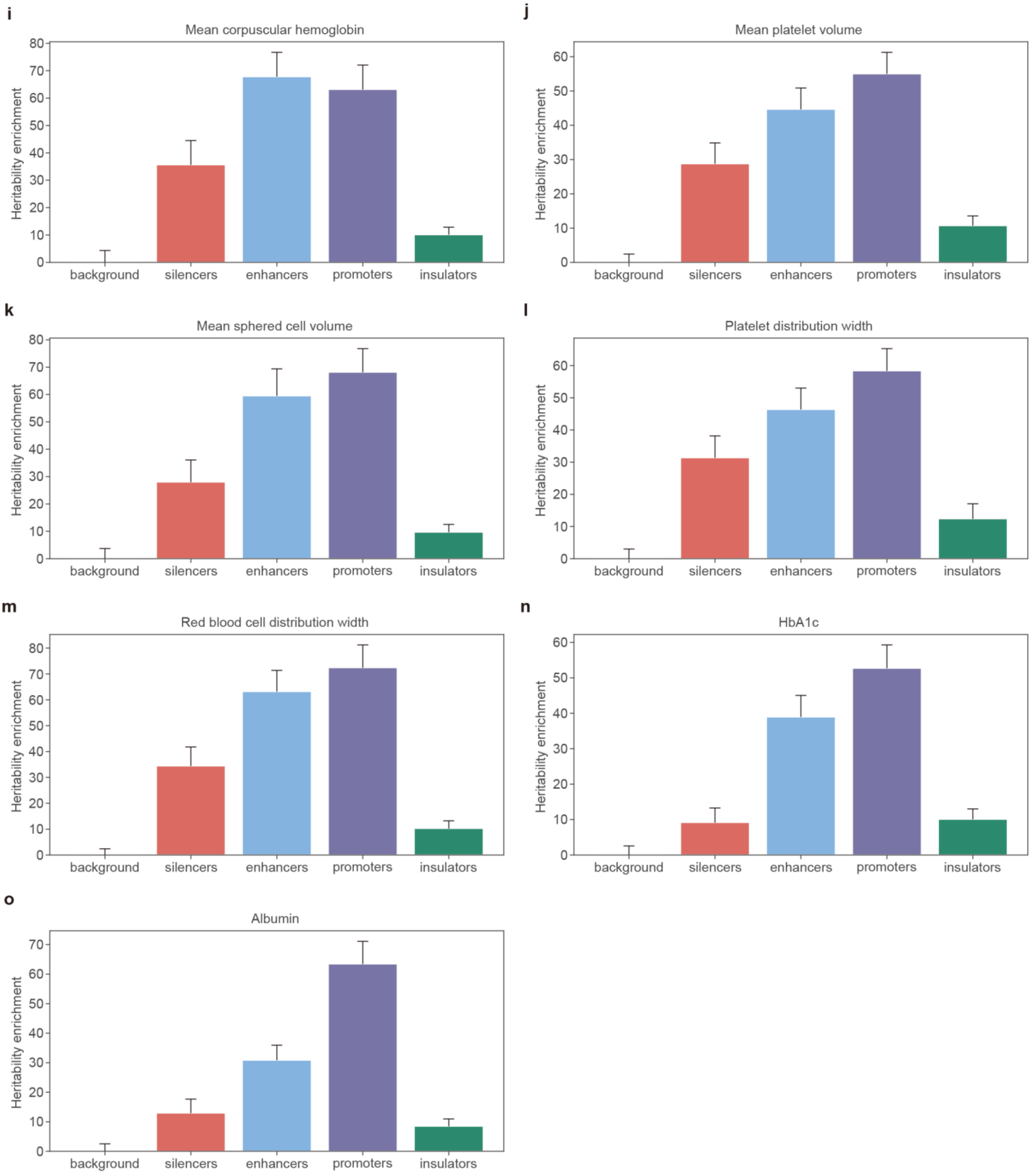

**Supplementary Fig. 18 (continue). Heritability enrichment analysis for true CREs and background regions on the K562 cell type. a-o**, Heritability enrichments estimated by LDSC within true CREs and background regions for blood-related traits including cancer (**a**), lymphocyte count (**b**), red count (**c**), white count (**d**), monocyte count (**e**), platelet count (**f**), eosinophil count (**g**), high light scatter reticulocyte count (**h**), mean corpuscular hemoglobin (**i**), mean platelet volume (**j**), mean sphered cell volume (**k**), platelet distribution width (**l**), red blood cell distribution width (**m**), HbA1c (**n**) and albumin (**o**). The error bars denote jackknife standard errors over 200 equally sized blocks of adjacent SNPs about the estimates of enrichment, and the centers of error bars represent the average value.

### Supplementary Tables

**Supplementary Table 1. Candidate CREs from different sources.**

| **Source** | **K562** | **HepG2** |
|---|---|---|
| DNase | 58,860 | 41,719 |
| ATAC | 61,509 | 45,112 |
| H2A.Z | 32,439 | 2880 |
| H3K9ac | 4677 | 5243 |
| H3K27ac | 8134 | 6359 |
| H3K4me1 | 28,320 | 24,647 |
| H3K4me2 | 13,416 | 15,270 |
| H3K4me3 | 7061 | 7488 |
| H3K9me1 | 2269 | - |
| H3K9me3 | 7036 | 48,882 |
| H3K27me3 | 36,780 | 32,722 |
| H3K36me3 | 3341 | 1343 |
| H3K79me2 | 1587 | 305 |
| H4K20me1 | 4830 | 486 |
| Total | 270,259 | 232,456 |
